# The impact of global and local Polynesian genetic ancestry on complex traits in Native Hawaiians

**DOI:** 10.1101/2020.05.18.102996

**Authors:** Hanxiao Sun, Meng Lin, Emily M. Russell, Ryan L. Minster, Tsz Fung Chan, Take Naseri, Muagututi‘a Sefuiva Reupena, Annette Lum-Jones, the Samoan Obesity, Lifestyle, and Genetic Adaptations (OLaGA) Study Group, Iona Cheng, Lynne R. Wilkens, Loïc Le Marchand, Christopher A. Haiman, Charleston W. K. Chiang

**Author notes:** These authors contributed equally (Hanxiao Sun, Meng Lin). Full membership listed in Supporting Information (the Samoan Obesity, Lifestyle, and Genetic Adaptations (OLaGA) Study Group).

## Abstract

Epidemiological studies of obesity, Type-2 diabetes (T2D), cardiovascular diseases and several common cancers have revealed an increased risk in Native Hawaiians compared to European- or Asian-Americans living in the Hawaiian islands. However, there remains a gap in our understanding of the genetic factors that affect the health of Native Hawaiians. To fill this gap, we studied the genetic risk factors at both the chromosomal and sub-chromosomal scales using genome-wide SNP array data on ∼4,000 Native Hawaiians from the Multiethnic Cohort. We estimated the genomic proportion of Native Hawaiian ancestry (“global ancestry,” which we presumed to be Polynesian in origin), as well as this ancestral component along each chromosome (“local ancestry”) and tested their respective association with binary and quantitative cardiometabolic traits. After attempting to adjust for non-genetic covariates evaluated through questionnaires, we found that per 10% increase in global Polynesian genetic ancestry, there is a respective 8.6%, and 11.0% increase in the odds of being diabetic (*P* = 1.65 10^−4^) and having heart failure (*P* = 2.18 10^−4^), as well as a 0.059 s.d. increase in BMI (*P* = 1.04 10^−10^). When testing the association of local Polynesian ancestry with risk of disease or biomarkers, we identified a chr6 region associated with T2D. This association was driven by an uniquely prevalent variant in Polynesian ancestry individuals. However, we could not replicate this finding in an independent Polynesian cohort from Samoa due to the small sample size of the replication cohort. In conclusion, we showed that Polynesian ancestry, which likely capture both genetic and lifestyle risk factors, is associated with an increased risk of obesity, Type-2 diabetes, and heart failure, and that larger cohorts of Polynesian ancestry individuals will be needed to replicate the putative association on chr6 with T2D.

**Author Summary:** Native Hawaiians are one of the fastest growing ethnic minority in the U.S., and exhibit increased risk for metabolic and cardiovascular diseases. However, they are generally understudied, especially from a genetic perspective. To fill this gap, we studied the association of Polynesian genetic ancestry, at genomic and subgenomic scale, with quantitative and binary traits in self-identified Native Hawaiians. We showed that Polynesian ancestry, which likely capture both genetic and non-genetic risk factors related to Native Hawaiian people and culture are associated with increased risk for obesity, type-2 diabetes, and heart failure. While we do not endorse utilizing genetic information to supplant current standards of defining community membership through self-identity or genealogical records, our results suggest future studies could identify population-specific genetic susceptibility factors that may be useful in suggesting underlying biological mechanisms and reducing the disparity in disease interventions in Polynesian populations.

## Introduction

Native Hawaiians are the second fastest growing ethnic group in the U.S., growing 40% from the 2000 to 2010 U.S. census [1]. Moreover, Native Hawaiians display alarming rates of obesity, coronary heart disease, diabetes, cardiovascular diseases, cancers, and other related chronic health conditions [2–9]. Epidemiological studies have shown that 49% of adult Native Hawaiians are obese, compared to 21% of European Americans and 13% of Japanese Americans living in Hawai‘i **Error! Reference source not found**., with > 2x and 5x higher odds of being obese than European- and Asian-Americans, respectively, after adjusting for socioeconomic status [6]. In addition, Native Hawaiians are ∼2-3 times more likely to develop Type-2 diabetes (T2D) than their European American counterparts, even after adjusting for common modifiable risk factors such as BMI and socioeconomic covariates [4]. Similarly, Native Hawaiians are ∼1.7 times more likely to develop cardiovascular diseases than European Americans [8], and cardiometabolic risk factors such as hypertension have been shown to be associated with genealogical estimates of proportion of Native Hawaiian ancestry [9]. Taken together, these observations suggest that in addition to non-genetic risk factors such as lifestyle or diet, there may be systematic differences in the number, frequency, or effect size of genetic risk alleles that contribute to epidemiological differences between Native Hawaiians and other continental populations. Yet, such genetic investigation has not been conducted and despite awareness and efforts to include more non-European populations in genomic studies, indigenous populations such as Native Hawaiians remain understudied [10–12].

Today, Native Hawaiians are an admixed population. Their ancestors settled the Hawai‘i archipelagos approximately 1,200-2,000 years ago and remained isolated there until 1778 when they encountered Western explorers who brought novel infectious agents that decimated the Native Hawaiian population before they rebounded over the last couple of centuries [13–16]. During the 18^th^ and 19^th^ centuries, Native Hawaiians became admixed with European and East Asian immigrants to the islands. The 2010 U.S. census data suggests that only approximately 1.2 million individuals in the U.S. derive some proportion of their ancestry from Native Hawaiians, accounting for about 0.4% of the U.S. population. The small population size may be one of the challenges in recruiting large cohorts, which contributes to the reason that this population is under-investigated from a genetic standpoint.

To begin filling the missing gap in the genetic understanding of disease risks in Native Hawaiians, we first distinguished a Native Hawaiian-specific component of ancestry from other continental ancestries, and tested the association of this global (genomic) ancestry to complex traits and diseases in Native Hawaiians. We presumed this component of ancestry to be Polynesian in origin, although we cannot discount the possibility that this component of ancestry has diverged from the prevalent ancestry component found in other extant Polynesian populations today. We further stress that associations between estimated global Polynesian ancestry and any phenotype will also capture any non-genetic cultural or environmental effects that are correlated with Polynesian ancestry. These variables are typically measured with considerable error; thus, adjustment for them does not exclude residual effects. Therefore, an observed association with genetic ancestry is not evidence for a deterministic impact attributed to the Polynesian genetic ancestry alone. Nevertheless, an observed association with genetic ancestry may imply that genetic mapping studies could identify genetic susceptibility factors enriched in the Polynesian populations that may be useful in suggesting underlying biological mechanisms.

We then tested the association of local Polynesian ancestry with complex traits and diseases in what is known as admixture mapping. Admixture mapping assumes that causal variants leading to increased risk or trait values occur more frequently on chromosomal segments inherited from the ancestral population that has higher disease risk or larger average trait values [17–19]. This technique is thus ideal as a first line analysis in understudied populations that are recently admixed. It has previously been used in African-American and Latino populations to identify novel genomic regions associated with phenotypes such as asthma, blood cell traits, breast and prostate cancer (reviewed in ref [17]), but has not yet been applied to Native Hawaiians.

## Results

### Impact of global genetic ancestry on cardiometabolic traits in Native Hawaiians

We used 3,940 self-identified Native Hawaiians from the Multiethnic Cohort (MEC) [20] that were genotyped on the MEGA array [21] to assess the impact of global ancestry on health. We first needed to construct a reference panel for Polynesian (PNS) ancestry since there is no publicly available reference panel for the PNS ancestry among Native Hawaiians. (Note: we refer to this ancestral component as Polynesian for simplicity.) Among the 3,940 Native Hawaiians in our dataset, we identified a panel of 178 unrelated Native Hawaiian individuals with the highest estimated amount of PNS ancestry (>90% in unsupervised ADMIXTURE analysis; **Methods**) after accounting for other sources of recent admixtures, namely Europeans (EUR), East Asians (EAS), and Africans (AFR). Using this reference panel, we computed a haplotype-based estimate of global genetic ancestry for each of the remaining 3,762 individuals, and kept 3,428 unrelated individuals after excluding for the first-degree relatedness in our dataset (**Methods**).

We then assessed in Native Hawaiians the association of each component of ancestries with a set of quantitative and binary cardiometabolic traits. Specifically, we focused on three disease categories for which the Native Hawaiians have shown increased risks in previous epidemiological studies: obesity [3,6], T2D [4], and cardiovascular disease [8,9]. We also examined quantitative traits and biomarkers associated with these diseases, namely BMI at baseline, fasting glucose and insulin level, HDL, LDL, triglycerides, and total cholesterol. More importantly, because non-genetic factors, such as socioeconomic status (SES) and lifestyle factors, could potentially confound the association between global genetic ancestry and risk of diseases, we attempted to adjust for these factors using education as individual level proxy to SES (**Methods**). Overall, we found that higher PNS ancestry is strongly associated with higher risk of obesity, T2D, heart failure (HF), and consistently, with higher BMI and lower HDL levels among the quantitative traits (**Table 1, S1-15 Tables**). For example, we observed that, holding the proportion of EAS and AFR ancestry constant, every 10% increase in the PNS ancestry in our cohort corresponded to a 0.059 s.d. (or 0.35 BMI unit) increase in BMI and a 1.09 times the odds of T2D (after adjusting for BMI). We observed opposite effects of PNS ancestry on waist-to-hip ratio (WHR) in males and females separately, though the statistical significance is marginal (**Table 1, S2 Table**). For T2D, HF, hypertension (HYPERT), and ischemic heart disease (IHD), BMI is an established risk factor. In our models, we also found BMI to be strongly associated with disease risk for these conditions (max *P* < 1 × 10^−7^; **S10-11, S13-14 Tables**). For T2D and HF, we observed a strong association between disease risk and PNS ancestry even after accounting for BMI, suggesting additional risk factors that are specific or correlated to the PNS ancestry (**Table 1**). For HYPERT and IHD, we observed a weak but nominally significant association between PNS ancestry and disease risk if we do not account for BMI. In fact, for most traits tested, the effect sizes due to PNS ancestry are lower after adjusting for BMI (**S1 Fig**), suggesting that at least part of the excessive risk for these traits may be mediated through BMI. Finally, other components of ancestry found in Native Hawaiians also exert an effect, as we observed that higher East Asian ancestry component are associated with increased risk of T2D, hyperlipidemia, and hypertension, but lower BMI and lowered risk of obesity (**Table 1**).

**Table 1:** summary of association between global genetic ancestry and quantitative and binary cardiometabolic traits in the Native Hawaiians.

| Trait | PNS |  | EAS |  | AFR |  |
| --- | --- | --- | --- | --- | --- | --- |
| | $\beta$ | P-value | $\beta$ | P-value | $\beta$ | P-value |
| Quantitative Traits |  |  |  |  |  |  |
| BMI | **0.5923** | **$1.04 \times 10^{-10}$** | **-0.6400** | **$<2 \times 10^{-16}$** | 1.0777 | 0.0948 |
| WHR (male) | -0.3592 | 0.0179 | -0.1358 | 0.2664 | 1.9487 | 0.1218 |
| WHR (female) | 0.2272 | 0.0985 | 0.2587 | 0.0139 | -0.1722 | 0.8524 |
| Glucose | -0.0129 | 0.929 | 0.1355 | 0.232 | 0.7825 | 0.463 |
| Insulin | 0.2858 | 0.0472 | 0.0048 | 0.966 | 0.6249 | 0.5573 |
| HDL | **-0.4715** | **$1.40 \times 10^{-4}$** | 0.1753 | 0.0700 | -1.4498 | 0.0988 |
| LDL | 0.0736 | 0.557 | 0.0720 | 0.463 | -0.5192 | 0.559 |
| TG | 0.1387 | 0.0342 | 0.1426 | 0.0053 | 0.1639 | 0.7237 |
| TC | -0.0441 | 0.7228 | 0.2072 | 0.0331 | -0.8084 | 0.3602 |
| Categorical Traits |  |  |  |  |  |  |
| Obesity | **1.2164** | **$2.24 \times 10^{-7}$** | **-1.3596** | **$5.40 \times 10^{-11}$** | 1.3181 | 0.3979 |
| T2D | **1.2416** | **$1.04 \times 10^{-9}$** | **0.6836** | **$2.05 \times 10^{-5}$** | 1.1393 | 0.4220 |
| T2D (adj BMI) | **0.8209** | **$1.65 \times 10^{-4}$** | **1.1765** | **$1.30 \times 10^{-11}$** | 0.3603 | 0.8125 |
| HF | **1.3104** | **$1.99 \times 10^{-6}$** | -0.0224 | 0.9209 | 3.4587 | 0.0493 |
| HF (adj BMI) | **1.0465** | **$2.18 \times 10^{-4}$** | 0.2528 | 0.2797 | 3.4653 | 0.0593 |
| HYPERL * | 0.0652 | 0.792 | **0.6973** | **$5.30 \times 10^{-4}$** | 0.9841 | 0.5731 |
| HYPERT | 0.6461 | 0.0100 | **0.7363** | **$2.74 \times 10^{-4}$** | -0.6385 | 0.7081 |
| HYPERT (adj BMI) | 0.3842 | 0.135 | **0.8783** | **$1.89 \times 10^{-5}$** | -0.9459 | 0.585 |
| IHD | 0.4787 | 0.0496 | 0.0164 | 0.9327 | -0.3750 | 0.8270 |
| IHD (adj BMI) | 0.2881 | 0.2457 | 0.1445 | 0.4633 | -0.7074 | 0.6871 |
| TIA | 0.4876 | 0.143 | -0.0201 | 0.941 | 2.3080 | 0.278 |
| TIA (adj BMI) | 0.3612 | 0.2839 | 0.0719 | 0.7928 | 2.1347 | 0.3226 |

We present the effect sizes ( $\beta$ , in units of s.d. for quantitative traits and log odds for binary traits) and p-values for the final model after accounting for covariates for PNS, EAS, and AFR ancestries, using proportion of EUR ancestry as baseline (**Method**). In all binary traits other than obesity, results adjusting for BMI as a
covariate in the model are also reported (\* BMI was not found to be associated with HYPERL and thus was not adjusted in the model). Effect sizes and P-values that are significant after adjusting for testing 14 traits are bolded. Abbreviations: BMI, body mass index; HDL, high-density lipoprotein; LDL, low-density lipoprotein; TG, triglycerides; TC, total cholesterol; T2D, Type-2 diabetes; HF, heart failure; HYPERL, hyperlipidemia; HYPERT, hypertension; IHD, ischemic heart disease; TIA, stroke and transient ischemic attack. For full model of each trait tested, please refer to **S1-15 Tables**.

Because, as mentioned above, non-genetic factors such as socioeconomic status could confound our analysis, we further tested if adding neighborhood SES could account for these associations. Neighborhood SES (nSES) is a validated composite measure created by principal component analysis that incorporates U.S. Census data on education, occupation, unemployment, household income, poverty, rent, and house values [22]. This nSES measure was categorized into quintiles based on the nSES distribution of Hawaii census tracts and Native Hawaiian subjects were assigned a quintile based on their geocoded baseline address (**Methods**). For BMI/obesity, HDL, T2D, and HF that showed significant association with proportion of PNS ancestry, adding nSES into the model showed that nSES was statistically significantly associated with each outcome, and accounted for some proportion of the risk. However, the association between proportion of PNS ancestry and each of these outcomes remained highly significant, with the exception of HDL, which became nominally significant (**Table 2, S1, S5, S9-11 Tables**). These results are again consistent with the possibility that unique Polynesian genetic risk factors exist in the Native Hawaiians that partly explain the elevated risk.

**Table 2:** summary of association between global genetic ancestry and quantitative and binary cardiometabolic traits in the Native Hawaiians, after adding nSES into the previous model that only adjusted for inidivdual level covariates.

|  | PNS |  | EAS |  | AFR |  |
| --- | --- | --- | --- | --- | --- | --- |
| Trait | $\beta$ | P-value | $\beta$ | P-value | $\beta$ | P-value |
| BMI | **0.4974** | **$2.26 \times 10^{-7}$** | **-0.6113** | **$1.37 \times 10^{-15}$** | 0.8537 | 0.201 |
| HDL | -0.3027 | 0.0201 | 0.1725 | 0.0891 | -0.9393 | 0.3021 |
| Obesity | **1.0774** | **$1.29 \times 10^{-5}$** | **-1.3105** | **$1.03 \times 10^{-9}$** | 1.2973 | 0.4141 |
| T2D (adj BMI) | **0.7575** | **$8.51 \times 10^{-4}$** | **1.1569** | **$6.92 \times 10^{-11}$** | 0.2535 | 0.8668 |
| HF (adj BMI) | **1.0201** | **$7.41 \times 10^{-4}$** | 0.2775 | 0.2588 | 3.4056 | 0.0714 |

We present the effect sizes ( $\beta$ , in units of s.d. for quantitative traits and log odds for binary traits) and p-values for the final model after accounting for covariates for PNS, EAS, and AFR ancestries, using proportion of EUR ancestry as baseline (**Method**). Effect sizes and P-values that are significant after adjusting for testing 14 traits are bolded.

As the strongest association with genetic ancestry came from BMI, we further investigated the association between BMI and PNS ancestry in stratified analysis. We found no evidence of difference between sexes (data not shown). However, we did observe a strong difference in the strength of association stratified by T2D disease status. Specifically, among T2D cases, we found no significant association between BMI and proportion of PNS ancestry (*P* = 0.112; **Fig 1, S16 Table**). On the other hand, among T2D controls, individuals were predicted to have 0.087 s.d. (or 0.51 units) or higher BMI per 10% increase in PNS ancestry (*P* = 1.4×10^−13^). This is despite the T2D strata having similar sample sizes (1,310 cases vs. 1,799 controls). A BMI model including interaction between T2D strata and PNS ancestry showed significant negative interaction (*P* = 0.0004, **S17 Table**). One interpretation is that relative individuals of other ancestries, BMI was only marginally increased among individuals with PNS ancestry when affected by T2D suggesting that there are alternative pathways (other than BMI) that contributes to T2D risk in the Polynesian population. This is consistent with our observation that PNS ancestry is independently associated with higher risk for T2D (**Table 1**, with adjustment of BMI).

**Fig 1:**
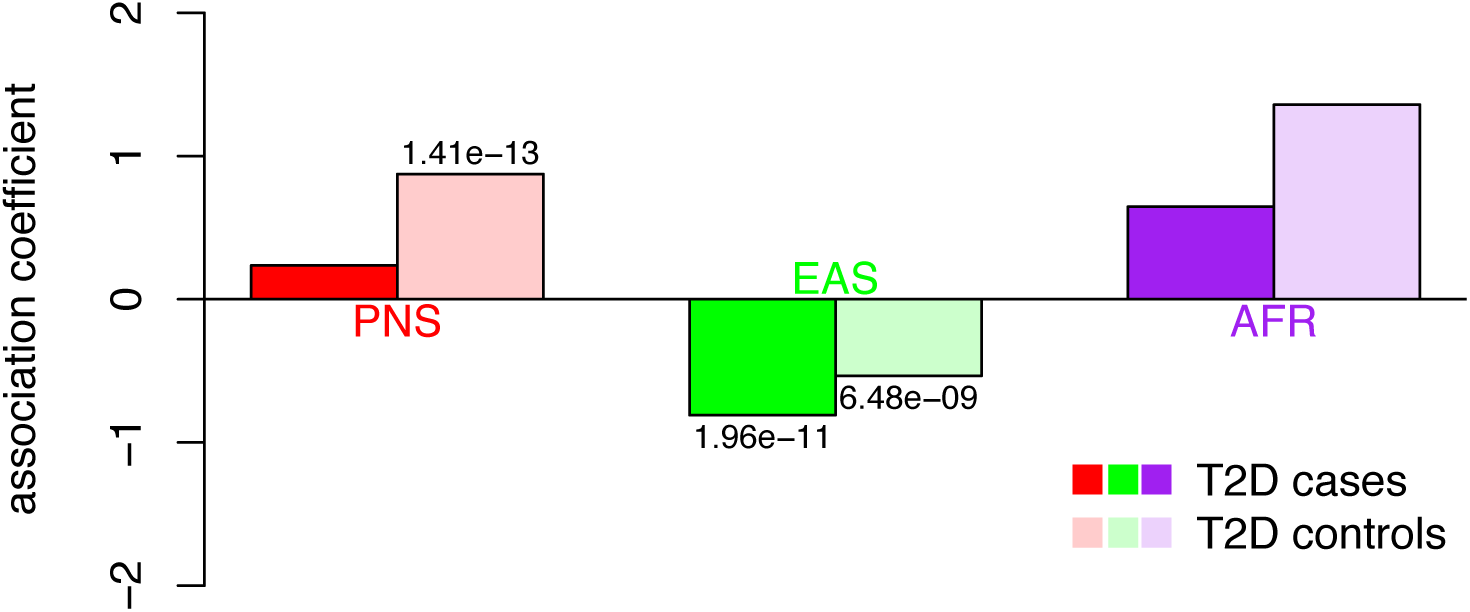
Stratified association testing between global genetic ancestry and BMI. Individuals were stratified based on T2D disease status. Cases are colored in darker color, controls in lighter color. P-values for significant association coefficients are provided. The strongly significant association between PNS ancestry and BMI among T2D controls, but not cases, is suggestive of an interaction between PNS ancestry and T2D.

For a subset of ∼300 Native Hawaiians in our cohort, we also have measures of subcutaneous fat and visceral fat, as well as lean mass vs. fat mass obtained through dual-energy x-ray absorptiometry and abdominal magnetic resonance imaging [23]. In this small subcohort, we found that increasing PNS ancestry to be more strongly and positively associated with subcutaneous fat (*P* = 4.88×10^−6^) compared to visceral fat (*P* = 0.014) (**Table 3**). There was no association with lean-to-fat mass ratio (*P* = 0.76), suggesting that PNS ancestry is associated with body fat distribution but not necessarily body fat composition. Because anthropometric measures of body fat distribution such as Waist-to-hip ratio often differ between male and females, we also conducted sex-stratified analysis. We observed similar trend of associations between subcutaneous fat vs. visceral fat, though the association seems more strongly driven by males (**Table 3**).

**Table 3:** Association of global ancestry with measures of fat distribution or fat composition among Native Hawaiians.

|  | Combined |  | Male |  | Female |  |
| --- | --- | --- | --- | --- | --- | --- |
|  | Estimate (s.e.) | P | Estimate (s.e.) | P | Estimate (s.e.) | P |
| Subcutaneous Fat |  |  |  |  |  |  |
| Intercept | -0.39 (0.25) | 0.11 | -0.58 (0.35) | 0.10 | -0.25 (0.37) | 0.51 |
| PNS | 1.30 (0.28) | **4.88x10<sup>-6</sup>** | 1.52 (0.4) | **2.29x10<sup>-4</sup>** | 1.14 (0.4) | **0.0052** |
| EAS | -0.14 (0.22) | 0.54 | 0.050 (0.33) | 0.88 | -0.28 (0.32) | 0.38 |
| AFR | 3.16 (1.80) | 0.080 | 5.45 (3.62) | 0.13 | 2.46 (2.12) | 0.25 |
| Total fat mass | -0.0044 (0.0081) | 0.59 | -0.0033 (0.013) | 0.80 | -0.0056 (0.011) | 0.62 |
| Visceral Fat |  |  |  |  |  |  |
| Intercept | -0.26 (0.26) | 0.31 | -0.11 (0.37) | 0.77 | -0.48 (0.38) | 0.20 |
| PNS | 0.71 (0.29) | **0.014** | 0.55 (0.42) | 0.19 | 0.98 (0.4) | **0.016** |
| EAS | -0.29 (0.23) | 0.21 | -0.092 (0.35) | 0.79 | -0.36 (0.32) | 0.27 |
| AFR | 3.45 (1.85) | 0.064 | 6.54 (3.82) | 0.089 | 2.45 (2.13) | 0.25 |
| Total fat mass | 0.0012 (0.0083) | 0.89 | -0.0082 (0.013) | 0.54 | 0.0073 (0.011) | 0.52 |
| Lean-to-Fat Mass Ratio |  |  |  |  |  |  |
| Intercept | 0.11 (0.18) | 0.55 | 0.49 (0.28) | 0.079 | -0.23 (0.25) | 0.37 |
| PNS | -0.087 (0.29) | 0.76 | -0.64 (0.42) | 0.13 | 0.40 (0.39) | 0.31 |
| EAS | -0.28 (0.24) | 0.24 | -0.77 (0.35) | 0.030 | 0.15 (0.33) | 0.64 |
| AFR | 0.86 (1.95) | 0.66 | 0.55 (4.21) | 0.90 | 1.02 (2.21) | 0.64 |

N = 280, 280, and 294 for analysis on subcutaneous fat, visceral fat, and lean-to-fat mass ratio, respectively. In sex-stratified analysis, N = 128, 128, and 136 males for analysis on subcutaneous fat, visceral fat, and lean-to-fat mass ratio, respectively; N = 152, 152, and 158 females for analysis on subcutaneous fat, visceral fat, and lean-to-fat mass ratio, respectively.

### Mapping of cardiometabolic traits using local genetic ancestry in Native Hawaiians

We next examined the impact of local genetic ancestry on cardiometabolic traits in Native Hawaiians through admixture mapping using linear or logistic regression models. We only analyzed the traits that exhibited a significant association with the global PNS ancestry. We used a threshold of 2.2×10^−5^ to declare genome-wide significance with a trait (**Method**).

Across the 2 quantitative (BMI and HDL) and 2 binary (T2D and HF; obesity was not included as the definition of obese status is dependent on BMI) traits examined through admixture mapping, we identified one region that surpassed our genome-wide significance threshold (**Fig 2**): 62.7Mb to 65.7Mb on chr6 for T2D (**Table 4, Fig 2**). We further defined a broader region encompassing neighboring regions with admixture *P*-value less than 1×10^−4^ as potential regions that may harbor causal allele(s). For this broader region spanning 11.4 Mb on chr6 (**Table 4**), we examined if known variants reported in the GWAS catalog could account for the signals we found through admixture mapping. We found 2 variants in the GWAS catalog for T2D that fall within our admixture peak (**S18 Table**). We imputed these two variants using 1000Genomes (phase 3) as the reference panel and found that conditioning on these variants did not significantly change our admixture mapping results (top P-value = 6.22×10^−6^; **S2 Figure**). These results suggest that our signals detected through admixture mapping may potentially be novel.

**Table 4:**
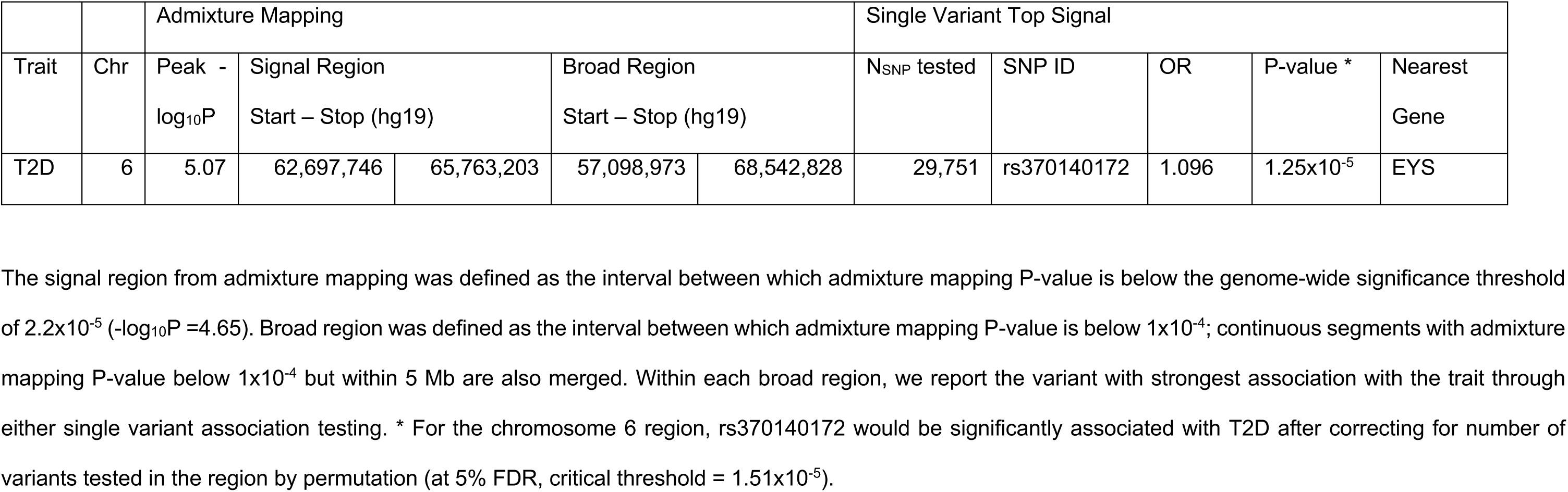
summary of significant loci identified through admixture mapping.

|  |  | Admixture Mapping |  |  |  |  | Single Variant Top Signal |  |  |  |  |
| --- | --- | --- | --- | --- | --- | --- | --- | --- | --- | --- | --- |
| Trait | Chr | Peak -<br>log<sub>10</sub>P | Signal Region<br>Start – Stop (hg19) |  | Broad Region<br>Start – Stop (hg19) |  | N<sub>SNP</sub> tested | SNP ID | OR | P-value * | Nearest<br>Gene |
| T2D | 6 | 5.07 | 62,697,746 | 65,763,203 | 57,098,973 | 68,542,828 | 29,751 | rs370140172 | 1.096 | 1.25x10<sup>-5</sup> | EYS |

The signal region from admixture mapping was defined as the interval between which admixture mapping P-value is below the genome-wide significance threshold of $2.2 \times 10^{-5}$ ( $-\log_{10}P = 4.65$ ). Broad region was defined as the interval between which admixture mapping P-value is below $1 \times 10^{-4}$ ; continuous segments with admixture mapping P-value below $1 \times 10^{-4}$ but within 5 Mb are also merged. Within each broad region, we report the variant with strongest association with the trait through either single variant association testing. \* For the chromosome 6 region, rs370140172 would be significantly associated with T2D after correcting for number of variants tested in the region by permutation (at 5% FDR, critical threshold = $1.51 \times 10^{-5}$ ).

**Fig 2:**
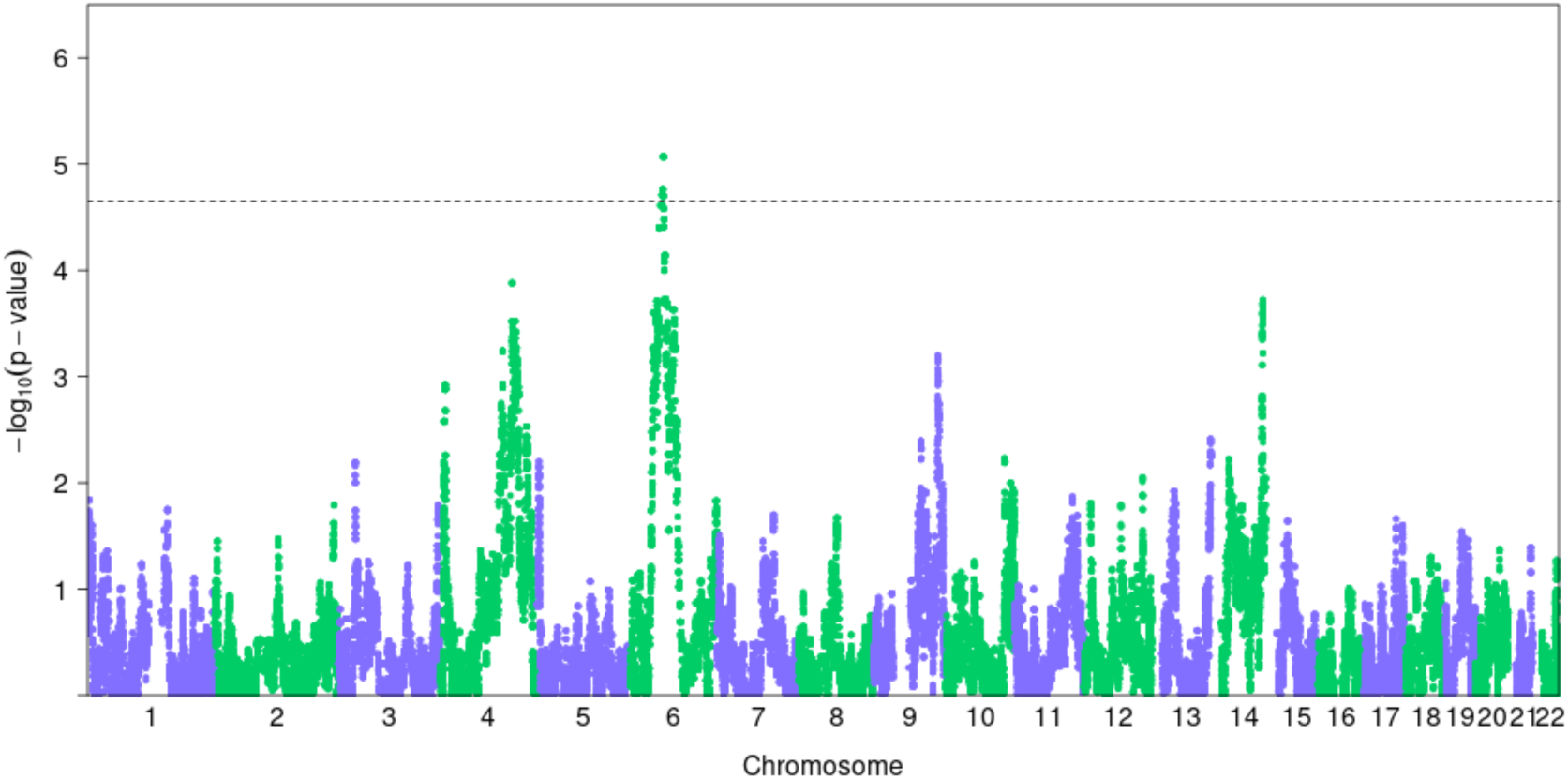
Manhattan plot of admixture mapping results for T2D. Dotted line denotes the genome-wide significance threshold for each trait at 2.2×10^−5^, determined through permutation.

To fine-map the candidate region on chromosome 6, we conducted single variant association tests (**Fig 3**). We imputed the full dataset of 3,940 individuals using 1000 Genomes as reference to increase coverage across the region, and accounted for cryptic relatedness and population structure in a logistic mixed model (**Methods**). We found that the top associated variant on chr6 for T2D was a well-imputed (INFO score = 0.86) 5’ UTR variant rs370140172 (OR = 1.096, *P* = 1.25×10^−5^, **Fig 3**). Its association with T2D was significant after accounting for number of markers tested in this region (regional significance threshold = 1.51×10^−5^; **Table 3**). This variant showed a large difference in frequency between Native Hawaiians (MAF = 24.2% among our reference PNS individuals; 11.2% among MEC-NH population) and European (0%) or East Asian (0.9%) individuals from 1000 Genomes (**S19 Table**). Conditioning on rs370140172 also drastically reduced the admixture association signal (minimum *P* ∼ 0.001 in the region; **S3 Fig**). Taken together, these observations suggest that rs370140172 or its proxy could be the allelic association driving the admixture signal.

**Fig 3.**
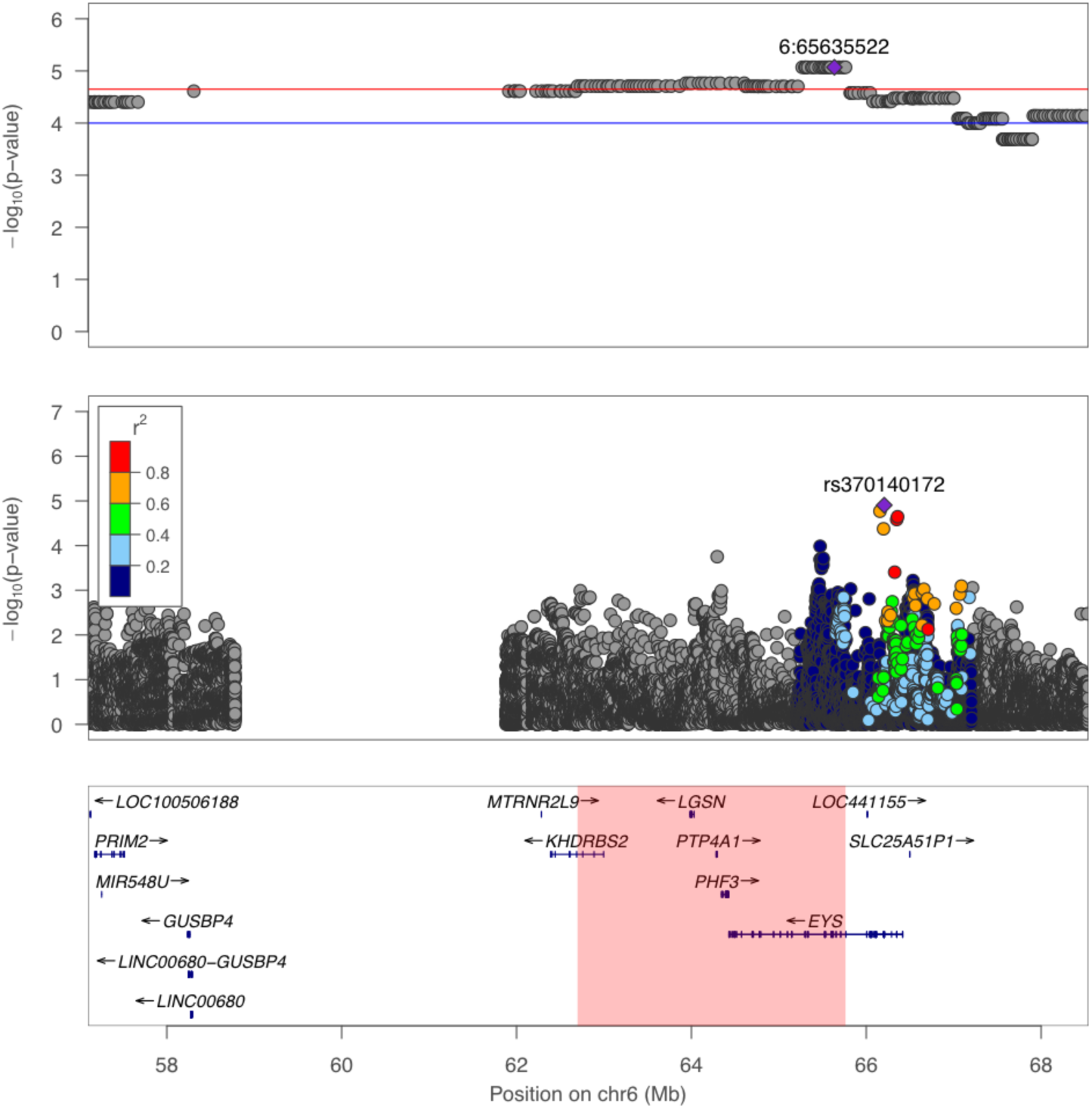
Association signals with T2D in the broad region of chr 6. The top panel depicts association signals from admixture mapping; middle panel depicts the single variation association, and bottom panel the genomic coordinates and nearby genes. Highlighted region at the bottom indicated the signal region as defined in Table 2.

We attempted to replicate the single variant signal on chromosome 6 by examining the lead variant and its proxies for association with T2D in another Polynesian population of 2,852 Samoan individuals, including 475 cases and 2,377 controls. We found no significant associations (minimum *P* = 0.743; **S20 Table**). However, we noted that the derived allele of rs370140172 had a significantly lower frequency in the Samoans (8.7%, compared to 24.2% in reference PNS individuals). The lower frequency of the allele and the current sample size does not provide sufficient power to replicate the association signal even at nominal significance level of 0.05 (Power = 14.2%; **S4 Figure**). Because of the difference in frequency between Native Hawaiians and Samoans, we also examined if this locus exhibits signals of positive natural selection in the Native Hawaiians (**Methods**). We found the derived allele to indeed sit on the longer haplotypes in Native Hawaiians, although the statistical significance is marginal compared to other loci of similar derived allele frequency in the genome (Z-score = +1.38; empirical *P* = 0.067). Thus, the elevated frequency in MEC-NH may still be the result of genetic drift.

## Discussion

This study aimed to fill a gap of genetic research in Native Hawaiians. We focused on studying the association of genetic ancestry, both globally and locally, to diseases for which Native Hawaiians showed increased risk. While the focus is on genetic ancestry, we emphasize that our approach does not constitute a methodology to quantify the degree of indigenousness among individuals native to the Hawaiian archipelago. Estimating proportion of genetic ancestry is not without errors, the results may change depending on the input genetic or reference data, and there is a conceptual difference between genetic ancestry and genealogical ancestry. Moreover, there are also difficulties in interpreting the estimated proportion; in this paper we made the simplifying assumption that the predominant component of ancestry found in MEC-NH individuals but not in other continental populations is Polynesian in origin. Given these caveats, we therefore believe the approach described here should not supplant current approaches, such as through self-reports or genealogical records, to define community membership. Consistent with this belief, we analyzed all individuals with available genetic data who self-identify as at least part Native Hawaiian ancestry; we did not attempt to define a population of Native Hawaiians using genetic data.

We began our analysis by modeling Polynesian ancestry. We first conducted ADMIXTURE analysis to identify an internal Native Hawaiian ancestry reference panel since there is no appropriate representative panel currently available. Consistent with their known history, we found Native Hawaiians to be a recently admixed population, deriving the largest proportion of their genetic ancestry from a presumed Polynesian ancestral component (on average ∼40.2%). We also found that global Polynesian ancestry from MEC-NH is positively and statistically significantly associated with BMI, HDL, Type-2 diabetes, obesity and heart failure after adjusting for other components of ancestries and available non-genetic covariates (**Table 1 and 2**). Polynesian ancestry was also nominally associated with WHR (in males), insulin level, triglycerides, hypertension and ischemic heart diseases, but these associations did not remain statistically significant after Bonferroni correction for the multiple traits that we tested in this study (**Table 1**). We then examined the association between local Polynesian ancestry and BMI, HDL, T2D, and HF. We found a 3.06 Mb region on chr6 possibly associated with Type-2 diabetes (**Fig 2** and **3**). Conservatively, we searched an expanded broader region encompassing 11.4Mb (**Fig 3, Table 4**) for known GWAS variants for T2D and showed that these known variants could not explain signal we detected (**S2 Fig, S18 Table**). Furthermore, single variant fine-mapping of the broad regions implicated a variant (rs370140172) on chr6 for T2D that was significantly associated with T2D after correcting for number of variants tested in permutation (**Table 4**) and showed large frequency differences between populations (**S19 Table**) that could account for the signal in local ancestry association (**S3 Fig**). Taken together, our findings suggest that these regions should be targeted for further investigation and replication in the future, preferably in additional Native Hawaiian or Polynesian populations.

The strong association between global ancestry and disease risks or related quantitative phenotypes suggests the presence of population-specific variants that could contribute to the increased risk observed in these populations. For example, a recently reported, Polynesian-specific, *CREBRF* variant discovered in Samoans was strongly associated with the odds of obesity, a finding that we previously replicated in Native Hawaiians [24]. However, we should also stress that an association with global ancestry would also in theory capture any non-genetic cultural or environmental effects that are correlated with ancestry. We attempted to control for non-genetic factors such as education, representing individual-level socioeconomic status (SES), and behavioral traits, such as cigarette smoking. We also examined the possibility that our observed associations with global ancestry was due to community-level SES by including neighborhood income levels in the model (**Table 2**). Admittedly, these variables are still imperfect proxies for SES and non-genetic factors certainly play a role in the etiology of these traits. Future studies may further integrate both individual-level (*e.g*. physical activity, diet, alcohol or medication use) and community-level (*e.g*. discrimination) non-genetic factors. Therefore we should interpret these associations with global ancestry with much caution.

These caveats notwithstanding, one notable observation is the association between global PNS ancestry and BMI. In analysis stratified by T2D status, despite having similar numbers of cases and controls, we found that PNS ancestry is not associated with BMI among T2D cases, but is associated with higher BMI among individuals unaffected by T2D (**S16 Table**). In models including interaction between global ancestry and T2D, we again observed that while T2D cases generally have higher BMI, those with greater PNS ancestry would actually have lowered BMI than those with less PNS ancestry (**S17 Table, Fig 1**). We interpret these findings to suggest that while PNS ancestry is positively associated with BMI, it is not proportionally increasing the risk of T2D compared to other ancestries. One possible explanation for this is through differential body composition. For example, individuals with increasing PNS ancestry may possess more lean mass, which contributes to BMI, than fat mass, which contributes to BMI and risk for T2D. There are some suggestions that individuals of PNS ancestry preferentially have greater lean mass than fat mass [25], although data is limited and we found no association between PNS ancestry and lean-to-fat mass ratio in a small subcohort of MEC-NH (**Table 2**). An alternative explanation is through differential fat distribution. For example, individuals with increasing PNS ancestry may preferentially store fat subcutaneously, which contribute to general adiposity and BMI but not necessarily to T2D, rather than viscerally, which could lead to insulin resistance and contribute to peripheral insulin sensitivity and further T2D [26,27]. We did find a stronger association between PNS ancestry with subcutaneous fat compared to visceral fat among our small sub-cohort of MEC-NH (**Table 2**), and it may be possible there are differences in deep versus superficial subcutaneous fat storage that we have not investigated. Ultimately, more data will be needed to make firm conclusions. What seems to be clear from our result is an independent pathway beyond BMI through which Native Hawaiians are also at risk for T2D. The negative interaction between T2D and PNS ancestry in the BMI model suggests individuals with increasing PNS ancestry are affected with T2D at lower BMI. In support of this hypothesis, we found that PNS ancestry is positively associated with risk of T2D even after adjusting for BMI (**Table 1**).

We conducted admixture mapping testing the association between local PNS ancestry genome-wide with the traits significantly associated with global PNS ancestry. Because admixture mapping had not been previously conducted among Native Hawaiians and the haplotypic pattern and LD structure within Native Hawaiians had not been previously explored, we used permutation to establish the genome-wide threshold for significance for a single trait, which we determined to be 2.2×10^−5^. Using this threshold, we found one notable region on chr6 associated with T2D.

Single variant fine-mapping of these regions showed a significant association on chr6 (rs370140172, P = 1.25×10^−5^) after correcting for number of variants tested by permutation (regional significance threshold = 1.51×10^−5^). This variant was imputed with high accuracy (INFO score = 0.86), exhibits large frequency enrichment compared to other populations (24% in non-admixed Native Hawaiians but monomorphic in 1KGP EUR and < 1% in 1KGP EAS; gnomAD v3 overall frequency = 0.00054), and explained the admixture mapping signal we detected (**S3 Fig**). Rs370140172 falls within the 5’ UTR (2^nd^ exon) of the gene *EYS*. Mutations in *EYS* can cause recessive retinitis pigmentosa [28,29], but there was no obvious link to T2D other than a suggestive association with T2D in Europeans (rs10498828, ∼670kb away, P=9×10^−6^), and a genome-wide association with BMI in a Japanese population (rs148546399, ∼1.5Mb away, P=1×10^−9^) [30,31]. Taken together, rs370140172 or its proxy may signal a novel population-specific candidate locus associated with T2D. We failed to replicate the association signal at this variant in a Samoan cohort. The failure to replicate could be partly explained by decreased power as the variant is rarer in Samoans and the cohort is relatively small. Furthermore, we may be limited by the availability of imputation panels; we used the 1000 Genome Project as reference panel and, as such, a number of Polynesian-specific variants that could underlie the admixture signal in these region may not be well imputed.

In summary, Native Hawaiians exhibit an increased risk for obesity, type-2 diabetes, and a number of cardiovascular diseases, but are generally understudied from a genetic standpoint in the literature. A better understanding of the genetic susceptibility risk factors will complement other epidemiological, non-genetic, risk factors for uniquely prevalent diseases among the Native Hawaiians. It is by integrating both genetic and non-genetic risk factors in our understanding of population-specific disease risk that we will have a better chance to control these diseases. Native Hawaiians have undergone a unique evolutionary history in their trans-Pacific voyages and settlement of the Hawaiian archipelago. Both the demographic and adaptive histories of these people may have shaped their genetic architecture. We present the first analysis of the genetic ancestry of present-day Native Hawaiians and suggest that it may have an impact on the risk of these diseases. However, genetic ancestry also reflects non-genetic cultural or environmental effects and we cannot exclude residual confounding by these variables. Nevertheless, if specific genetic susceptibility variants could be identified, they may be useful in clarifying underlying biological mechanisms. Further studies focusing on indigenous Polynesian populations, such as Native Hawaiians, will advance the findings reported here and may help alleviate the disparity in genomic medical research existing for Native Hawaiians.

## Materials and Methods

### Study population

In this study, we used genetic and epidemiologic data from Native Hawaiian individuals from the Multiethnic Cohort (MEC). MEC is a prospective epidemiological cohort of >215,000 individuals spanning five major ethnicities, including biospecimen samples on >5,300 Native Hawaiians. It is currently the largest single cohort with genetic information on Native Hawaiians, and thus is ideal for our study. In this study, we used a subcohort of >3,900 individuals genotyped on the MEGA genotyping array [21] as part of the PAGE consortium [32]. The institutional review boards of the University of Hawai‘i and the University of Southern California approved the study protocol. All participants signed an informed consent form.

Quality control of MEGA array was previously described [24]. In general, individual and genotype level quality control filters were previously applied as part of PAGE, and additionally we applied the following steps: All variant names were updated to dbSNP v144; duplicated loci and indels were removed; triallelic variants or variants with non-matching alleles to 1000 Genomes Project phase 3 (1KGP) were discarded; loci with unique positions not found in 1KGP were removed from the dataset; alleles were standardized to the positive strand by comparing to 1KGP. Finally, a genotype missingness filter of 5% and a minor allele frequency filter of 1% were applied, resulting in a total of 3,940 MEC Native Hawaiian (MEC-NH) individuals genotyped at 697,505 SNPs.

### Global and local ancestry inference

In addition to a predominant Polynesian (PNS) ancestry, Native Hawaiians are known to be recently admixed with individuals of European and East Asian ancestry [14]. In order to define individual genetic ancestry, whether locally or globally, we needed a reference panel for the Polynesian component of the Native Hawaiian ancestry. As such a reference panel does not exist, we sought to construct an internal reference panel by identifying MEC-NH individuals with the largest amount of global Polynesian ancestry as previously described [24]. Briefly, we combined all MEC individuals genotyped on the same MEGA array (3,940 Native Hawaiians, 3,465 Japanese, 30 Hispanic/Latinos, 5,325 African Americans) and all individuals from 1000 genomes Project, pruned SNPs with r^2^ > 0.1 (using window sizes of 50 SNPs with steps of 10 SNPs across the genome), and partitioned the samples to two groups of related (up to and including 2^nd^ degree) and unrelated individuals by KING (default threshold used). We then ran ADMIXTURE (v. 1.3.0) in unsupervised mode for unrelated samples, then projected the estimated ancestral allele frequency to the related samples to infer the genomic ancestries of the related group. We found stable estimates at k=4 after 5 iterations. MEC-NH individuals at k=4 exhibited known components of ancestry from European, East Asian and African, as well as a component of ancestry that is unique to the MEC-NH, presumed to be Polynesian (**S5 Fig**). We then identified 178 unrelated MEC-NH individuals (kinship coefficient <0.2 estimated from PC-relate [33]) whose Polynesian component of ancestry were estimated to be over 0.9 as reference for the Polynesian component of the Native Hawaiian ancestry.

To call local ancestry, we merged the MEC-NH samples with the above 1000 Genomes reference individuals, and rephrased the merged dataset using EAGLE2 (v 2.4.1). Next, we combined the 178 MEC-NH reference with the above 1KGP reference individuals to form the reference panel. Using this reference panel, we then inferred local ancestry used RFMix [34] (version 2.03-r0). One key parameter for RFMix is the local recombination rates, which vary across continental populations [35,36] but has not been estimated for Native Hawaiians or Polynesians. However, using multi-way admixed 1KGP American (AMR) populations, we evaluated the impact of misspecification of a recombination map. We found that RFMix inferences of local ancestry are robust even using a constant recombination map (>98% concordance, **S21 Table**). Therefore we used HapMap2 pooled recombination map (ftp://ftp-trace.ncbi.nih.gov/1000genomes/ftp/technical/working/20110106_recombination_hotspots/) to infer local ancestry in Native Hawaiians. To obtain global ancestry estimates, we summed the local ancestry estimates across the genome, after excluding tracts that have any ancestral probability < 0.9. We observed that on average, a self-reported Native Hawaiian individual derived ∼29.6% ancestry from EUR, ∼29.0% ancestry from EAS, ∼1.2% ancestry from AFR, and the remaining ∼40.2% ancestry from PNS. These values are similar to previous estimates of proportions of genetic ancestry from MEC using ancestry informative markers [37]. The summed PNS ancestry from RFMix is highly concordant with that inferred from ADMIXTURE [24], and is thus used for phenotype association and covariate adjustments in admixture mapping (below).

### Phenotype transformation

We focused on three categories of traits for which the Native Hawaiians exhibit excess risk in past epidemiological studies [2–4,6–9]: (1) adiposity traits, which include BMI at baseline and obesity; (2) metabolic traits, which include fasting glucose level, fasting insulin level, and Type-2 diabetes (T2D); and (3) cardiovascular traits, which include HDL, LDL, triglycerides (TG), total cholesterol (TC), heart failure (HF), hyperlipidemia (HYPERL), hypertension (HYPERT), ischemic heart disease (IHD), and stroke and transient ischemic attacks (TIA). Fasting glucose and insulin levels were collected after entry to the MEC, between 2001-2006. Obesity, T2D, HF, HYPERL, HYPERT, IHD, and TIA are binary disease outcomes. The metabolic and the quantitative cardiovascular traits were previously studied by PAGE consortium; we thus followed the inclusion criteria and phenotype transformation (based on medication use) as previously suggested by PAGE [32] (**S22 Table**). At a given BMI, Polynesians have a higher proportion of lean muscle mass to fat mass than Europeans so we use the recommended BMI cut-off of 32 kg/m^2^ to define obesity cases and controls [25,38]. T2D includes prevalent cases at cohort entry and incident cases during follow-up, based on self-report with medication use in questionnaires or a report from linkage to Hawai‘i insurers, CMS, or CHDD [32]. For incident binary cardiovascular traits we utilized the Medicare fee-for-service linkage data for MEC [39] defined as https://www2.ccwdata.org/web/guest/condition-categories. Descriptive summaries of the traits and covariates can be found in **S23 Table**.

### Associations between binary and quantitative traits with global ancestries

We tested the association of global Polynesian ancestry with quantitative and binary traits using linear and logistic regressions, respectively. We focused on the 3428 unrelated individuals after removing first degree relatives determined by KING [40] and individuals used in the internal PNS reference panel. To account for the impact of non-genetic factors that can confound the association between traits and genetic ancestry, covariate-adjusted outcomes were create by regressing out the impact of the non-genetic factors. These include behavioral traits such as smoking and education, as proxies for socioeconomic status. For quantitative traits, we first conducted univariate regression of the trait of interest on the non-genetic covariates. We then retained age and sex in the model, as well as all covariates that are nominally significantly associated with the trait. For categorical covariates retained in this procedure, we grouped the non-significantly associated level to reduce the variable down to a ternary or binary variable. We then model the covariates jointly in a multivariate regression model, and then standarized the residuals from this model. The standardized residuals were then used in a multivariate regression model with estimated global Polynesian, East Asian and African ancestries as independent variables, leaving European as the reference. For binary traits, we maintained the same structure, first removing uncorrelated covariates based on univariate logistic regression models. The remaining covariates are then used in a multivariate logistic regression with the addition of global ancestry estimates. The coefficients and p-values associated with the non-genetic covariates and global ancestries from the multivariate regression model are provided in **S1-15 Tables**.

### Adjusting for neighborhood socioeconomic status (nSES) measures

To further assess if the association between global ancestry and outcome could be explained by uncaptured non-genetic factors, we included the nSES variable in our regression models [22]. We determined nSES by subjects’ residential census tract using an index derived from principal components of indicator variables of SES (education level; proportion unemployed and with blue collar job; proportion <200% poverty line; proportion employed; median household income, rent and home value) based on 1990 Hawaii Census data. Each Native Hawaiian geocoded baseline address (1993-1996) was assigned a nSES qunitle based on the distribution of neighborhood SES across all census tracts in Hawaii. For traits that showed strong association in **Table 1** (*i.e*. BMI/obesity, HDL, T2D, and HF), we added nSES in the model to account for confounding in the assessment of the association with global ancestry. Because of the area-based design, Native Hawaiian participants residing in the same census tract were assigned the same nSES measure. We thus used a mixed effect model to account for this spatial clustering by including the census tract ID as random effect. We used *lmer* and *glmer* (version 1.1-21) function in R (version 3.6.2) with default parameters.

### Mapping of binary and quantitative traits using local Polynesian ancestry

To identify local genomic segments in which the Polynesian ancestry is associated with a trait of interest, we conducted admixture mapping using logistic or linear regression. We focused on the same 3428 unrelated individuals used in global ancestry analysis (above). We used linear or logistic regression to test the association of estimated dosage of Polynesian ancestry from RFMix at each genomic location, while controlling for estimated global ancestry from EUR, EAS, and AFR. Traits were modeled in the same way as above in the global ancestry analysis, except we focused only on individual-level covariates for computational efficiency of genome-wide testing.

We determined the significance threshold for admixture mapping for a given trait using two approaches: by a recently published simulation-based approach [41] and by permutation. For the simulation-based approach, because we were only interested in testing the association of Polynesian ancestry to a trait of interest, we dichotomized estimated local ancestry into Polynesian and non-Polynesian segments to estimate the covariance in local ancestry across the genome. We then estimated the genome-wide significance threshold in admixture mapping to be 2.28×10^−5^ using 10,000 simulations in STEAM [41]. For permutation-based approach, we simulated 1,000 runs of genome-wide admixture mapping, each based on a random phenotype drawn from a standard normal distribution. We then examined the distribution of the most significantly associated p-value from each of the simulations and set at the 5% false discovery level to the threshold of 2.24×10^−5^. The two thresholds are nearly identical, and are similar to previously suggested threshold among Latinos [42] (4.8×10^−5^). We thus used 2.2×10^−5^ as the genome-wide significance threshold for admixture mapping, and also considered regions with local ancestry association p-values between 1×10^−4^ and 2.2×10^−5^ as suggestive and report these findings.

### Conditional analysis and single variant tests in associated admixture region

For the locus we identified through local ancestry association (**Table 4**), we defined the signal region as contiguous variants with admixture *P*-values lower than the genome-wide significance threshold (2.2×10^−5^, or -log_10_P>4.64). We then defined a broad region by extending the signal region to nearby flanking regions that are (1) < 5Mbp away upstream or downstream from the signal region, and (2) with -log_10_P>4. We then imputed our rephased dataset using Sanger Imputation Service (https://imputation.sanger.ac.uk/). We used 1KGP as the reference panel, and PBWT as the imputation software. We subsequently filtered out indels and loci with low imputation quality (INFO score <0.4), and applied a minor allele frequency filter of 1%. We then investigated whether a previously known variant from the GWAS catalog [43] for the same trait could drive this signal by including all GWAS catalog variants residing in the broad region and passed quality control in our study as covariates in a conditional regression analysis.

We also conducted single variant association based on imputed dosages in the entire broad region. We included all 3,940 samples in this analysis, and corrected the relatedness by using a linear mixed model from EMMAX [44]. The inter-sample relatedness was calculated from PC-relate [33] so to be freed from possible population structure. We followed the same covariate model and phenotype transformation as was done in admixture mapping, except for using the top 10 principal components (PCs) from PC-air [45] as substitutes for the global ancestry covariates. 1,000 permutations were carried out to estimate the regional critical values for significance.

### Replication analysis in Samoans

We attempted to replicate the association of rs370140172 and nine other proxies showing the strongest single-variant associations with a cross-sectional population based study of Samoans recruited from Independent Samoa in 2010 [38,46]. This study was approved by the institutional review board of Brown University and the Health Research Committee of the Samoa Ministry of Health. All participants gave written informed consent via consent forms in Samoan language.

The Samoan participants from 2010 were genotyped genome-wide with Affymetrix 6.0 genotyping arrays [38]. A subset of 1,284 Samoan participants were whole-genome sequenced as part of the Trans-Omics for Precision Medicine (TOPMed) Program sponsored by the National Institutes of Health (NIH) National Heart, Lung, and Blood Institute (NHLBI). The sequences were used to produce a Samoan-specific reference panel for genotype imputation. Genotypes absent from the Affymetrix genotyping array and present in the reference panel were imputed in the remaining Samoan participants. T2D case and control exclusion criteria were defined to mirror that used in the MEC-NH analyses. Specifically, we removed cases who were pregnant, diagnosed with type 1 diabetes, or under 20 years old. We removed controls with fasting glucose greater than 7 mmol/L. This resulted in 475 cases and 2,377 controls. Association testing was conducted using logistic mixed model regression implemented in lme4qtl [47]. Empirical kinship as estimated from the genotypes was included as a random effect covariate. Age, BMI, education (coded as a continuous variable in six levels), and the first ten PCs were included as fixed effect covariates in the logistic mixed model regression.

The Power of the replication analysis was conducted using the Genetic Association Study power calculator (http://csg.sph.umich.edu/abecasis/cats/gas_power_calculator/), assuming the case control sample size, estimated frequency of rs370140172, and a prevalence rate of T2D of 17.1% [46] in Samoans, and a significance threshold of 0.05.

### Test of Natural Selection

We calculated the nSL score [48] of derived alleles across all imputed loci using Selscan [49], after the post imputation quality control. We calculated nSL among 178 MEC-NH reference individuals who had estimated PNS ancestry > 90%, and compared the nSL value for rs370140172 (derived allele of 0.24, and INFO score of 0.87) to that of 44,266 variants selected from the genome matched by imputation uncertainty (INFO score 0.77-0.97) and derived allele frequency (0.23-0.25).

## Acknowledgements

We would like to thank all of the Native Hawaiian participants in the Multiethnic Cohort that are involved in this study, which was funded through grants from the National Cancer Institute (U01CA164973, P01CA168530) and National Human Genome Research Institute (U01HG007397). We would also like to thank University of Hawai‘i Cancer Center’s Native Hawaiian Community Advisory Board for reviewing the study proposal and providing comments to earlier version of this manuscript. Furthermore, we would like to thank the Samoan participants of the study, local village authorities, and the many Samoan and other field workers who contributed to the success of the Samoan OLaGA Study Group. We acknowledge the Samoan Ministry of Health and the Samoa Bureau of Statistics for their support of this research. Whole genome sequencing (WGS) for the Trans-Omics in Precision Medicine (TOPMed) program was supported by the National Heart, Lung and Blood Institute (NHLBI). NHLBI TOPMed: Genome-wide Association Study of Adiposity in Samoans was funded by NHLBI (R01HL093093 [S.T. McGarvey] and (R01 HL133040 [R.L.M.]); WGS (phs000972) was performed at the University of Washington Northwest Genomics Center (HHSN268201100037C) and the New York Genome Center (HHSN268201500016C). Centralized read mapping and genotype calling, along with variant quality metrics and filtering were provided by the TOPMed Informatics Research Center (3R01HL-117626-02S1; contract HHSN268201800002I). Phenotype harmonization, data management, sample-identity QC, and general study coordination were provided by the TOPMed Data Coordinating Center (3R01HL-120393-02S1; contract HHSN268201800001I). We gratefully acknowledge the studies and participants who provided biological samples and data for TOPMed.

## Supporting Information

**S1 Fig: Correlation of effect sizes attributed to PNS ancestry in the regression model with or without adjustment for BMI**. Across the binary traits tested, even if the effect attributable to PNS ancestry is not significant, the effect sizes are lowered if accounting for BMI, suggesting at least part of the excess risk for these traits among Native Hawaiians are mediated through higher BMI associated with the ancestry. Hyperlipidemia was excluded because BMI is not associated with the disease risk in univariate regression model. HF, heart failure; HYPERT, hypertension; IHD, ischemic heart disease; T2D, type-2 diabetes; TIA, stroke and transient ischemic attack.

**S2 Fig: Admixture mapping P-value with or without conditioning on variants previously reported in GWAS catalog to be associated with T2D (rs79976124 and rs10498828)**. Green and blue colors denote SNP level P-value in association testing with and without, respectively, conditioning on known GWAS variants.

**S3 Fig: Admixture mapping P-value after conditioning on the most strongly associated variant in single variant analysis in chr6 (T2D) broad region**. The originally reported admixture signal (blue) can be explained by the conditioned variant (green), suggesting that these single variants might be novel variants associated with these traits.

**S4 Fig: Power of replicating the top signal from single variant analysis with T2D in Samoans**. We estimated the power to replicate the top signal (rs370140172) from single variant analysis with T2D in the Samoan cohort, using GAS power calculator (http://csg.sph.umich.edu/abecasis/cats/gas_power_calculator/index.html). The prevalence rate of T2D in Samoans set as 17.1%, which was the value averaged over the reported values in both sex. The number of cases (N=475) and controls (N=2377) were set to the observed sample size in Samoans. The genotype relative risk was set to estimated OR (1.096) from MEC-NH.

**S5 Fig: Global ancestry proportion estimated from unsupervised ADMIXTURE analysis, after integrating runs of relatedness and unrelatedness**. 3,465 MEC Japanese (MEC-JA), 30 MEC Latinos (MEC-LA), 5,325 MEC African Americans (MEC-AA), and 3,940 MEC Native Hawaiians (MEC-NH) were merged with the 1000 Genomes Project populations. At K = 4 we identified an ancestral component (colored red) that are found largely in Native Hawaiians, presumed to be the Polynesian ancestry.

**S1 Table: Details of the association statistics of the covariates and global ancestries of BMI**. Model 1 models the non-genetic covariates according to the heuristic described in the **Methods**. The residual from model 1 is then inverse normalized and tested in model 2. Models 1A and 2A repeats the procedure but included quintiles of nSES levels in a mixed effect model (**Method**); in this case, the R^2^ in Model 1A reported include both the fixed and the random effect. * edu4 was a binary variable created from the original categorical variable of education status by grouping levels 1,2,3 and coded 0, while education status level 4 was coded as 1. This was done because there were no significant associations between education levels 1 through 3 and BMI. See Supplemental Table 21 for description of these education levels.

**S2 Table: Details of the association statistics of the covariates and global ancestries of WHR**. Model 1 models the non-genetic covariates according to the heuristic described in the **Methods**. The residual from model 1 is then inverse normalized and tested in model 2. The top panels were conducted in males only; the bottom in females only. See Supplemental Table 21 for description of these education and cigarette smoking levels.

**S3 Table: Details of the association statistics of the covariates and global ancestries of fasting glucose**. Model 1 models the non-genetic covariates according to the heuristic described in the **Methods**. The residual from model 1 is then inverse normalized and tested in model 2.

**S4 Table: Details of the association statistics of the covariates and global ancestries of fasting insulin**. Model 1 models the non-genetic covariates according to the heuristic described in the **Methods**. The residual from model 1 is then inverse normalized and tested in model 2.

**S5 Table: Details of the association statistics of the covariates and global ancestries of HDL**. Model 1 models the non-genetic covariates according to the heuristic described in the Methods. The residual from model 1 is then inverse normalized and tested in model 2. Models 1A and 2A repeats the procedure but included quintiles of nSES levels in a mixed effect model (Method); in this case, the R^2^ in Model 1A reported include both the fixed and the random effect. **S6 Table: Details of the association statistics of the covariates and global ancestries of LDL**. Model 1 models the non-genetic covariates according to the heuristic described in the Methods. The residual from model 1 is then inverse normalized and tested in model 2.

**S7 Table: Details of the association statistics of the covariates and global ancestries of TG**. Model 1 models the non-genetic covariates according to the heuristic described in the Methods. The residual from model 1 is then inverse normalized and tested in model 2. * edu4 was a binary variable created from the original categorical variable of education status by grouping levels 1,2,3 and coded 0, while education status level 4 was coded as 1. This was done because there were no significant associations between education levels 1 through 3 and BMI.

**S8 Table: Details of the association statistics of the covariates and global ancestries of total cholesterol**. Model 1 models the non-genetic covariates according to the heuristic described in the **Methods**. The residual from model 1 is then inverse normalized and tested in model 2.

**S9 Table: Details of the association statistics of the covariates and global ancestries of obesity**. Model 1 models the non-genetic covariates according to the heuristic described in the Methods. Model 2 then includes global ancestries in addition to the significant covariates. * edu4 was a binary variable created from the original categorical variable of education status by grouping levels 1,2,3 and coded 0, while education status level 4 was coded as 1. This was done because there were no significant associations between education levels 1 through 3 and obesity. Model 3 included quintiles of nSES levels in a mixed effect model.

**S10 Table: Details of the association statistics of the covariates and global ancestries of Type-2 Diabetes**. Model 1 models the non-genetic covariates according to the heuristic described in the **Methods**. Model 2 then includes global ancestries in addition to the significant covariates. Model 3 included quintiles of nSES levels in a mixed effect model. * edu3 was a ternary variable created from the original categorical variable of education status by grouping levels 1 and 2. This was done because there were no significant associations between education levels 1 and 2 with T2D.

**S11 Table: Details of the association statistics of the covariates and global ancestries of heart failure**. Model 1 models the non-genetic covariates according to the heuristic described in the Methods. Model 2 then includes global ancestries in addition to the significant covariates. Model 3 included quintiles of nSES levels in a mixed effect model. * edu3 was a ternary variable created from the original categorical variable of education status by grouping levels 1 and 2. This was done because there were no significant associations between education levels 1 and 2 with heart failure.

**S12 Table: Details of the association statistics of the covariates and global ancestries of hyperlipidemia**. Model 1 models the non-genetic covariates according to the heuristic described in the Methods. Model 2 then includes global ancestries in addition to the significant covariates. * edu4 was a binary variable created from the original categorical variable of education status by grouping levels 1,2,3 and coded 0, while education status level 4 was coded as 1. This was done because there were no significant associations between education levels 1 through 3 and hyperlipidemia.

**S13 Table: Details of the association statistics of the covariates and global ancestries of hypertension**. Model 1 models the non-genetic covariates according to the heuristic described in the **Methods**. Model 2 then includes global ancestries in addition to the significant covariates. **S14 Table: Details of the association statistics of the covariates and global ancestries for ischemic heart disease**. Model 1 models the non-genetic covariates according to the heuristic described in the Methods. Model 2 then includes global ancestries in addition to the significant covariates. * edu3 was a ternary variable created from the original categorical variable of education status by grouping levels 1 and 2. This was done because there were no significant associations between education levels 1 and 2 with ischemic heart disease.

**S15 Table: Details of the association statistics of the covariates and global ancestries for stroke and transient ischemic attacks**. Model 1 models the non-genetic covariates according to the heuristic described in the **Methods**. Model 2 then includes global ancestries in addition to the significant covariates.

**S16 Table: Stratified analysis of association between global genetic ancestry and BMI among T2D cases and controls**. Model testing was performed in the same manner as the global analysis with BMI (**S1 Table**), except for stratifying based on T2D disease status. Model column provided the final model with association coefficients. * edu4 was a binary variable created from the original categorical variable of education status by grouping levels 1,2,3 and coded 0, while education status level 4 was coded as 1. This was done because there were no significant associations between education levels 1 through 3 and BMI.

**S17 Table: Model of association between global ancestry and BMI, including interaction with type-2 diabetes**. Model 1 models the non-genetic covariates according to the heuristic described in the Methods, except for type-2 diabetes status. The residual from model 1 was then inverse normalized and tested in model 2, which includes global ancestries, type-2 diabetes status, and interactions between global ancestries and type-2 diabetes status. * edu4 was a binary variable created from the original categorical variable of education status by grouping levels 1,2,3 and coded 0, while education status level 4 was coded as 1. This was done because there were no significant associations between education levels 1 through 3 and BMI.

**S18 Table: Variants within the admixture signal region that were reported to be associated with the tested or related traits in GWAS catalog**. Reported P-value, associated trait, and mapped genes were provided by the GWAS catalog. Allele frequencies were either calculated from the imputed data of the 178 reference MEC Native Hawaiian individuals with estimated PNS ancestry > 90%, or obtained from 1000 Genomes Project. Frequencies were reported with respect to the minor allele in the Native Hawaiians, given in parenthesis next to the Native Hawaiian frequency estimates.

**S19 Table: Allele frequencies across populations for the most strongly associated variant in chr6 for T2D in single variant association test**. Allele frequencies were either calculated from the imputed data of the 178 reference MEC Native Hawaiian individuals with estimated PNS ancestry > 90%, or obtained from 1000 Genomes Project (reported on dbSNP: https://www.ncbi.nlm.nih.gov/snp/). Frequencies were reported with respect to the derived allele, given in parenthesis next to the Native Hawaiian frequency estimates.

**S20 Table: Association results to T2D in 2**,**852 Samoan Replication Cohort**. We attempted to replicate the association of rs370140172 and nine other proxies showing the strongest single-variant associations with a cross-sectional population based study of Samoans recruited from Independent Samoa (**Methods**). EAF, effect allele frequency in Samoans. BETA and SE refers to the effect size and standard errors, respectively, from the logistic mixed model association tests in the Samoan cohort. P-val (Samoa) and P-val (MEC-NH) provide the p-value from the logistic mixed model association tests in the Samoan cohort and MEC Native Hawaiian cohort, respectively.

**S21 Table: Local ancestry inference using RFMix is robust to the choice of recombination map**. To evaluate the impact of recombination map on local ancestry inference, we used the 1000 Genomes AMR population. Following the same procedure used for Native Hawaiians, we identified through unsupervised ADMIXTURE analysis 49 Peruvian (PEL) and 3 Mexican (MEX) individuals from 1000 Genomes as having > 80% Native American ancestry. We then inferred local ancestry using RFMix in 71 HapMap3 MEX individuals using the constructed reference panel of 99 CEU, 108 YRI, and 52 NA individuals from 1000 Genomes. We used three recombination map in the local ancestry inference: a HapMap2 pooled recombination map, a mis-specified African-American map, and a constant map that assumes a constant rate of 1cM / Mb across the genome. We compared in pairwise fashion the concordance of inferred ancestry across common variants between runs, and calculated concordance rate as the sum of the diagonal of the contingency table. Across all comparisons, even when using a constant rate map, the concordance rate is extremely high (0.987, 0.981, and 0.981 for the comparisons of default vs. AA map, default to constant rate map, and constant rate to AA map, respectively), suggesting that the choice of recombination map does not strongly impact the local ancestry inference using RFMix.

**S22 Table: phenotype inclusion and transformation for metabolic and quantitative cardiovascular traits**. These traits were studied in PAGE consortium and we thus follow the same criteria and transformation.

**S23 Table: Descriptive summary statistics of the traits and covariates analyzed**. Summary statistics reported after exclusion and transformation as described in S20 Table. For biomarkers (glucose, insulin, HDL, LDL, TG, and TC), a subset of participants were invited after cohort entry. Thus there is an age at baseline and an age at blood draw.

