## Supplementary Information for "The impact of global and local Polynesian genetic ancestry on complex traits in Native Hawaiians"

**Supporting Information**

Additional members of the OLaGA Study Group includes:

Jenna C. Carlson[8], Jerry Z. Zhang[8], Nicola L. Hawley[9], Rajan Deka[10], Daniel E. Weeks[2,8], Stephen T. McGarvey[11,12]

[8]Department of Biostatistics, Graduate School of Public Health, University of Pittsburgh, Pittsburgh, PA
[9]Department of Epidemiology (Chronic Disease), Yale School of Public Health, New Haven, Connecticut, USA
[10]Department of Environmental Health, College of Medicine, University of Cincinnati, Cincinnati, Ohio, USA
[11]International Health Institute and Department of Epidemiology, School of Public Health, Brown University, Providence, Rhode Island, USA
[12]Department of Anthropology, Brown University, Providence, Rhode Island, USA

**Supplemental Figures**

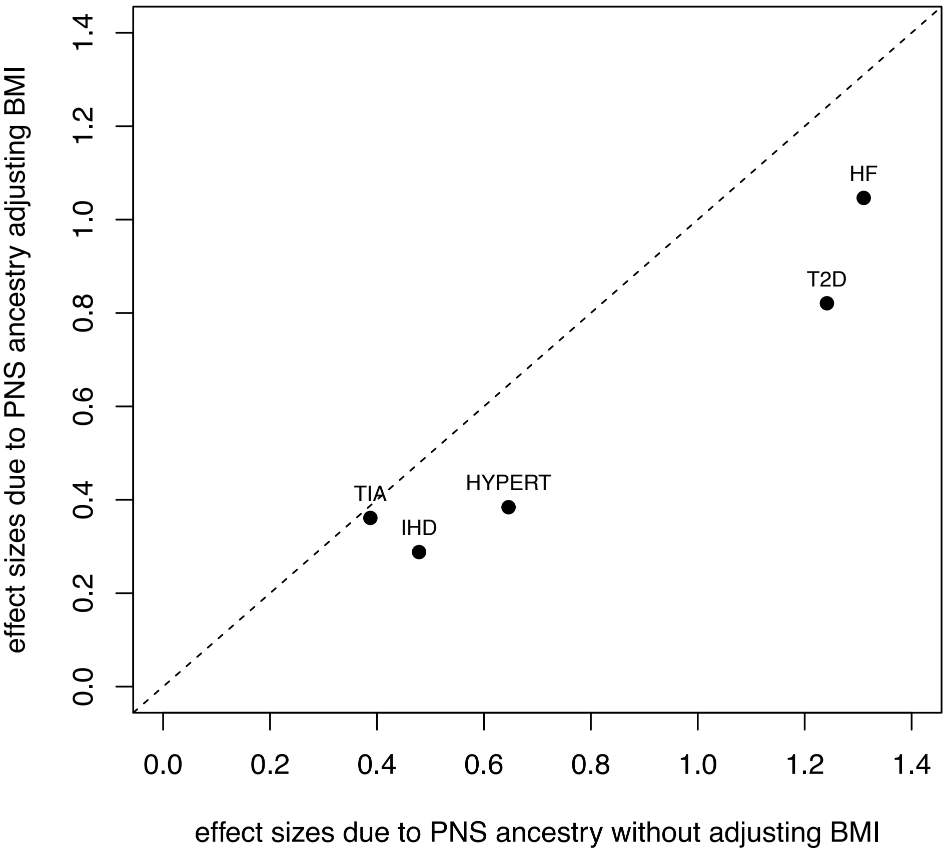


**S1 Fig: Correlation of effect sizes attributed to PNS ancestry in the regression model with or without adjustment for BMI.** Across the binary traits tested, even if the effect attributable to PNS ancestry is not significant, the effect sizes are lowered if accounting for BMI, suggesting at least part of the excess risk for these traits among Native Hawaiians are mediated through higher BMI associated with the ancestry. Hyperlipidemia was excluded because BMI is not associated with the disease risk in univariate regression model. HF, heart failure; HYPERT, hypertension; IHD, ischemic heart disease; T2D, type-2 diabetes; TIA, stroke and transient ischemic attack.

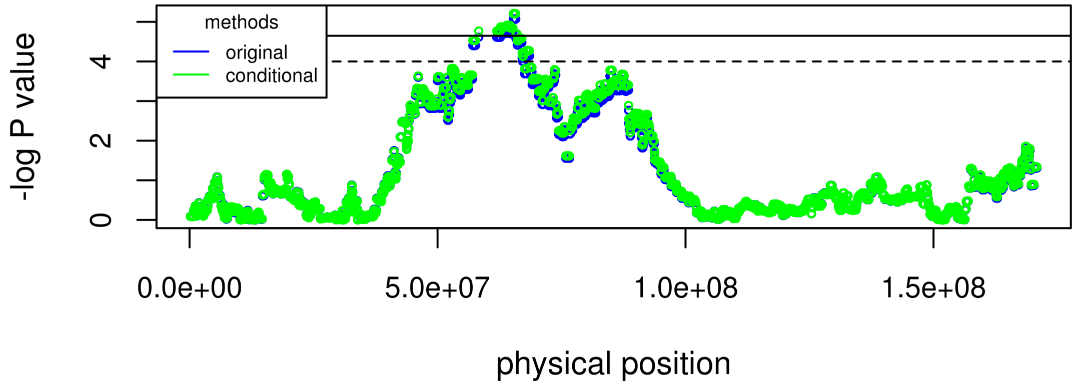


**S2 Fig: Admixture mapping P-value with or without conditioning on variants previously reported in GWAS catalog to be associated with T2D (rs79976124 and rs10498828)**. Green and blue colors denote SNP level P-value in association testing with and without, respectively, conditioning on known GWAS variants.

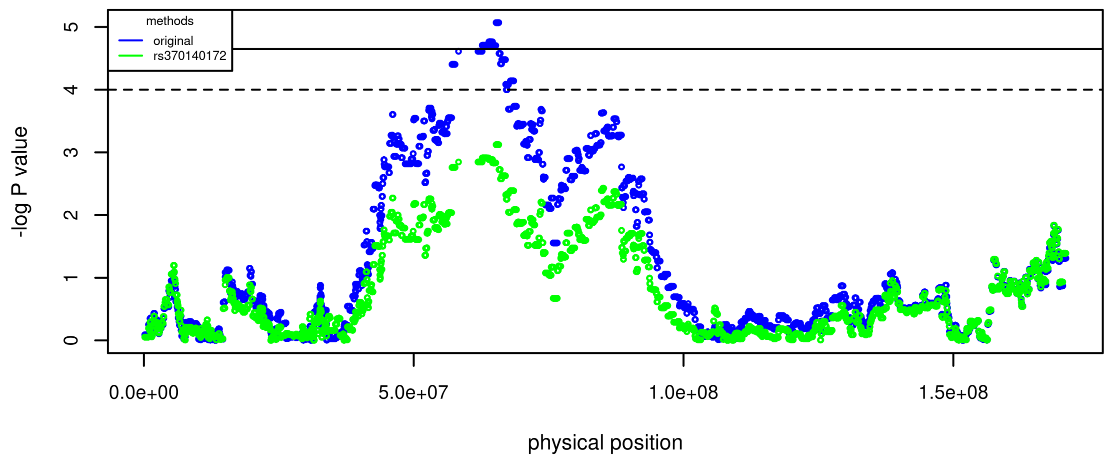


**S3 Fig: Admixture mapping P-value after conditioning on the most strongly associated variant in single variant analysis in chr6 (T2D) broad region**. The originally reported admixture signal (blue) can be explained by the conditioned variant (green), suggesting that these single variants might be novel variants associated with these traits.

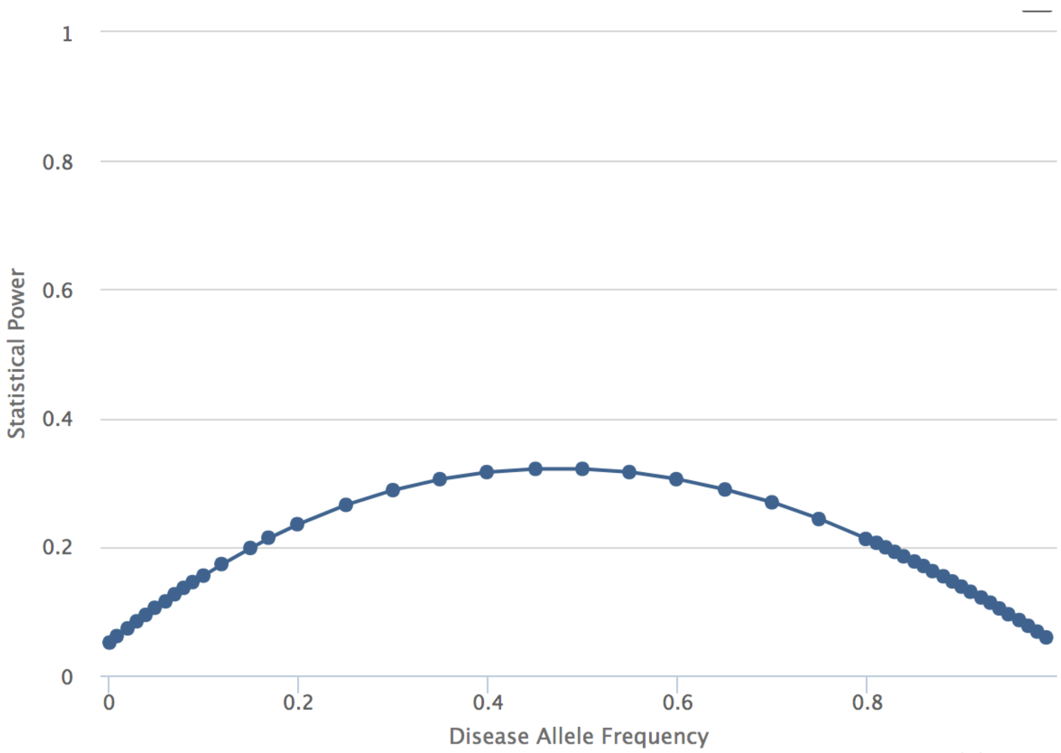


**S4 Fig: Power of replicating the top signal from single variant analysis with T2D in Samoans.** We estimated the power to replicate the top signal (rs370140172) from single variant analysis with T2D in the Samoan cohort, using GAS power calculator (http://csg.sph.umich.edu/abecasis/cats/gas_power_calculator/index.html). The prevalence rate of T2D in Samoans set as 17.1%, which was the value averaged over the reported values in both sex [1]. The number of cases (N=475) and controls (N=2377) were set to the observed sample size in Samoans. The genotype relative risk was set to estimated OR (1.096) from MEC-NH.

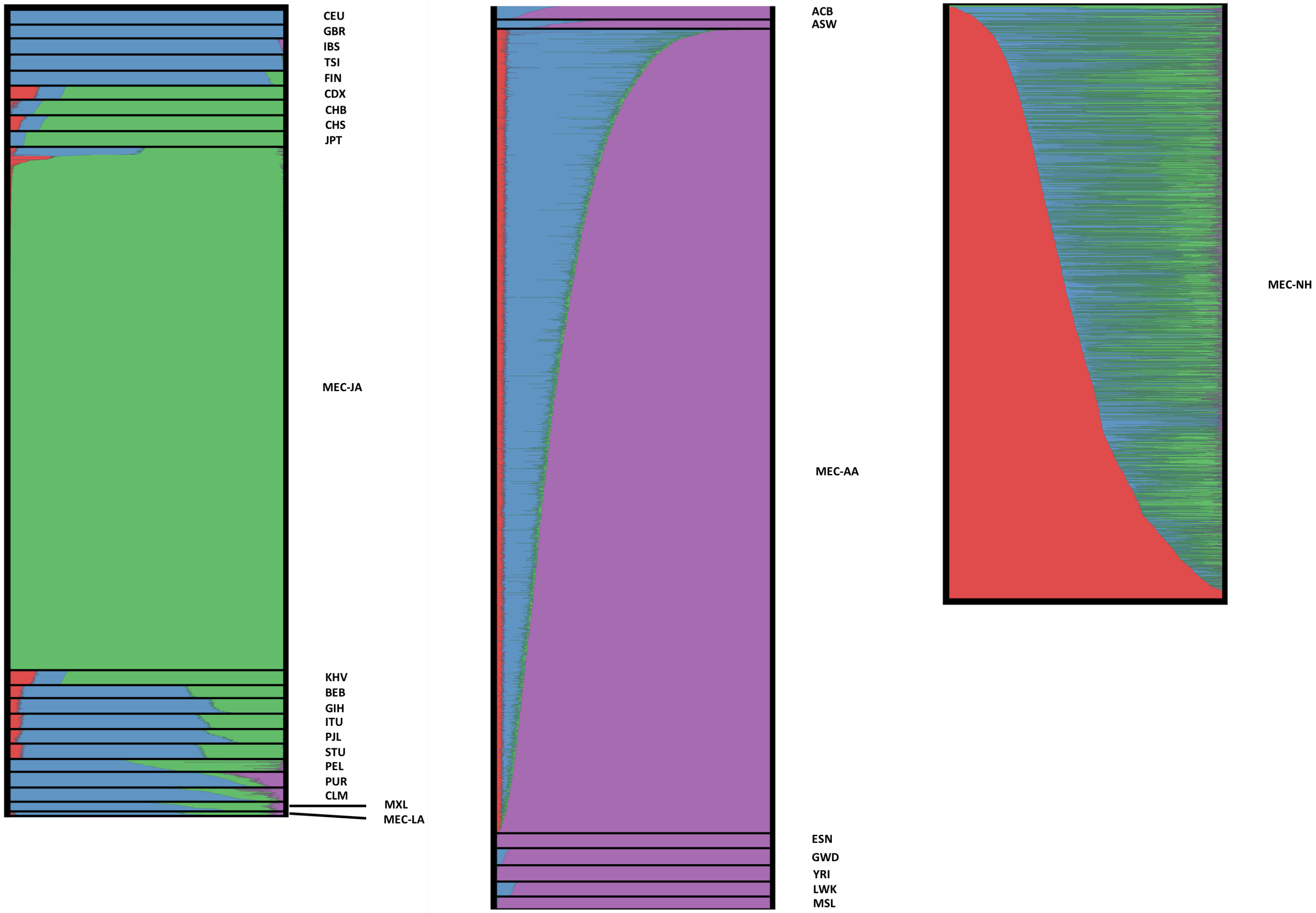


**S5 Fig: Global ancestry proportion estimated from unsupervised ADMIXTURE analysis, after integrating runs of relatedness and unrelatedness.** 3,465 MEC Japanese (MEC-JA), 30 MEC Latinos (MEC-LA), 5,325 MEC African Americans (MEC-AA), and 3,940 MEC Native Hawaiians (MEC-NH) were merged with the 1000 Genomes Project populations. At K = 4 we identified an ancestral component (colored red) that are found largely in Native Hawaiians, presumed to be the Polynesian ancestry.

**Supplemental Tables**

### S1 Table: Details of the association statistics of the covariates and global ancestries of BMI.

<table>
<tr><td colspan="8">Model 1: linear regression between BMI and covariates</td></tr>
<tr><th colspan="2">variables</th><th>estimate</th><th>std. error</th><th>t</th><th>p</th><th>$R^2$</th><th>df</th></tr>
<tr><td colspan="2">intercept</td><td>30.7091</td><td>0.9639</td><td>31.861</td><td>$<2\times10^{-16}$</td><td rowspan="7">0.1449</td><td rowspan="7">3081</td></tr>
<tr><td colspan="2">age (at baseline)</td><td>-0.0418</td><td>0.0176</td><td>-2.383</td><td>0.0173</td></tr>
<tr><td colspan="2">t2d</td><td>13.6203</td><td>1.4981</td><td>9.092</td><td>$<2\times10^{-16}$</td></tr>
<tr><td colspan="2">sex</td><td>-1.3251</td><td>0.2611</td><td>-5.074</td><td>$4.12\times10^{-7}$</td></tr>
<tr><td colspan="2">edu4*</td><td>-1.0763</td><td>0.2319</td><td>-4.641</td><td>$3.62\times10^{-6}$</td></tr>
<tr><td colspan="2">t2d:age</td><td>-0.1925</td><td>0.0266</td><td>-7.225</td><td>$6.28\times10^{-13}$</td></tr>
<tr><td colspan="2">t2d:sex</td><td>1.5213</td><td>0.4014</td><td>3.79</td><td>$1.54\times10^{-4}$</td></tr>
<tr><td colspan="8">Model 2: linear regression between standardized residual and global ancestry</td></tr>
<tr><td colspan="2">intercept</td><td>-0.0638</td><td>0.0543</td><td>-1.175</td><td>0.2402</td><td rowspan="4">0.0625</td><td rowspan="4">3084</td></tr>
<tr><td colspan="2">PNS</td><td>0.5923</td><td>0.0914</td><td>6.484</td><td>$1.04\times10^{-10}$</td></tr>
<tr><td colspan="2">EAS</td><td>-0.6400</td><td>0.0726</td><td>-8.817</td><td>$<2\times10^{-16}$</td></tr>
<tr><td colspan="2">AFR</td><td>1.0777</td><td>0.6449</td><td>1.671</td><td>0.0948</td></tr>
</table>

<table>
<tr><td colspan="8">Model 1A: linear mixed model between BMI and covariates, including nSES</td></tr>
<tr><td colspan="2">intercept</td><td>31.9467</td><td>1.0524</td><td>30.355</td><td>$<2\times10^{-16}$</td><td rowspan="11">0.1574</td><td rowspan="11">2840</td></tr>
<tr><td colspan="2">age (at baseline)</td><td>-0.0445</td><td>0.0185</td><td>-2.399</td><td>0.0165</td></tr>
<tr><td colspan="2">t2d</td><td>12.9035</td><td>1.5541</td><td>8.303</td><td>$<2\times10^{-16}$</td></tr>
<tr><td colspan="2">sex</td><td>-1.4258</td><td>0.2766</td><td>-5.155</td><td>$2.72\times10^{-7}$</td></tr>
<tr><td colspan="2">edu4*</td><td>-0.6812</td><td>0.2483</td><td>-2.743</td><td>0.0061</td></tr>
<tr><td rowspan="4">nSES</td><td>(Q2 vs. Q1)</td><td>-0.5086</td><td>0.4023</td><td>-1.264</td><td>0.2082</td></tr>
<tr><td>(Q3 vs. Q1)</td><td>-1.4301</td><td>0.3862</td><td>-3.703</td><td>$2.95\times10^{-4}$</td></tr>
<tr><td>(Q4 vs. Q1)</td><td>-0.9863</td><td>0.3835</td><td>-2.572</td><td>0.0112</td></tr>
<tr><td>(Q5 vs. Q1)</td><td>-1.7015</td><td>0.3709</td><td>-4.587</td><td>$1.00\times10^{-5}$</td></tr>
<tr><td colspan="2">t2d:age</td><td>-0.1821</td><td>0.0276</td><td>-6.606</td><td>$4.69\times10^{-11}$</td></tr>
<tr><td colspan="2">t2d:sex</td><td>1.4948</td><td>0.4171</td><td>3.584</td><td>$3.45\times10^{-4}$</td></tr>
<tr><td colspan="8">Model 2A: linear regression between standardized residual, including nSES, and global ancestry</td></tr>
<tr><td colspan="2">intercept</td><td>-0.0331</td><td>0.0569</td><td>-0.583</td><td>0.56</td><td rowspan="4">0.0516</td><td rowspan="4">2836</td></tr>
<tr><td colspan="2">NH</td><td>0.4974</td><td>0.0958</td><td>5.189</td><td>$2.26\times10^{-7}$</td></tr>
<tr><td colspan="2">EAS</td><td>-0.6113</td><td>0.0761</td><td>-8.034</td><td>$1.37\times10^{-15}$</td></tr>
<tr><td colspan="2">AFR</td><td>0.8537</td><td>0.6681</td><td>1.278</td><td>0.201</td></tr>
</table>

Model 1 models the non-genetic covariates according to the heuristic described in the **Methods**. The residual from model 1 is then inverse normalized and tested in model 2. Models 1A and 2A repeats the procedure but included quintiles of nSES levels in a mixed effect model (**Method**); in this case, the $R^2$ in Model 1A reported include both the fixed and the random effect. * edu4 was a binary variable created from the original categorical variable of education status by grouping levels 1,2,3 and coded 0, while education status level 4 was coded as 1. This was done because there were no significant associations between education levels 1 through 3 and BMI. See Supplemental Table 21 for description of these education levels.

**S2 Table: Details of the association statistics of the covariates and global ancestries of WHR**.

| Model 1: linear regression between WHR and covariates in males | | | | | | | |
|---|---|---|---|---|---|---|---|
| variables | | estimate | std. error | t | p | $R^2$ | df |
| intercept | | 0.8655 | 0.0207 | 41.88 | $<2\times10^{-16}$ | 0.1126 | 1150 |
| age | | -0.0002 | 0.0002 | -0.859 | 0.3905 | | |
| bmi | | 0.0037 | 0.0004 | 10.386 | $<2\times10^{-16}$ | | |
| edu | (2 vs. 1) | -0.0063 | 0.0080 | -0.784 | 0.4334 | | |
| | (3 vs. 1) | -0.0097 | 0.0080 | -1.203 | 0.2291 | | |
| | (4 vs. 1) | -0.0192 | 0.0083 | -2.321 | 0.0205 | | |
| cig | (1&2&3 vs. 4) | 0.0130 | 0.0042 | 3.071 | 0.0022 | | |
| | (1&2&3 vs. 5) | 0.0150 | 0.0051 | 2.929 | 0.0035 | | |
| | (1&2&3 vs. 6) | 0.0128 | 0.0071 | 1.796 | 0.0728 | | |
| Model 2: linear regression between standardized residual and global ancestry in males | | | | | | | |
| intercept | | 0.1613 | 0.0923 | 1.748 | 0.0808 | 0.0089 | 1155 |
| PNS | | -0.3592 | 0.1515 | -2.372 | 0.0179 | | |
| EAS | | -0.1358 | 0.1221 | -1.112 | 0.2664 | | |
| AFR | | 1.9487 | 1.2585 | 1.548 | 0.1218 | | |

| Model 1: linear regression between WHR and covariates in females | | | | | | | |
|---|---|---|---|---|---|---|---|
| variables | | estimate | std. error | t | p | $R^2$ | df |
| intercept | | 0.7832 | 0.0204 | 38.347 | $<2\times10^{-16}$ | 0.0638 | 1494 |
| age | | 0.0005 | 0.0003 | 1.964 | 0.0497 | | |
| bmi | | 0.0028 | 0.0003 | 8.597 | $<2\times10^{-16}$ | | |
| edu | (2 vs. 1) | -0.0272 | 0.0089 | -3.041 | 0.0024 | | |
| | (3 vs. 1) | -0.0231 | 0.0090 | -2.556 | 0.0107 | | |
| | (4 vs. 1) | -0.0296 | 0.0094 | -3.153 | 0.0017 | | |
| cig | (1&2&3 vs. 4) | 0.0083 | 0.0052 | 1.595 | 0.1108 | | |
| | (1&2&3 vs. 5) | 0.0192 | 0.0076 | 2.517 | 0.0120 | | |
| | (1&2&3 vs. 6) | 0.0048 | 0.0122 | 0.394 | 0.6934 | | |
| Model 2: linear regression between standardized residual and global ancestry in males | | | | | | | |
| intercept | | -0.1606 | 0.0780 | -2.06 | 0.0395 | 0.0050 | 1499 |
| PNS | | 0.2272 | 0.1374 | 1.653 | 0.0985 | | |
| EAS | | 0.2587 | 0.1051 | 2.462 | 0.0139 | | |
| AFR | | -0.1722 | 0.9257 | -0.186 | 0.8524 | | |

Model 1 models the non-genetic covariates according to the heuristic described in the **Methods**. The residual from model 1 is then inverse normalized and tested in model 2. The top panels were conducted in males only; the bottom in females only. See Supplemental Table 21 for description of these education and cigarette smoking levels.

**S3 Table: Details of the association statistics of the covariates and global ancestries of fasting glucose.**

| Model 1: linear regression between glucose and covariates | | | | | | |
|---|---|---|---|---|---|---|
| variables | estimate | std. error | t | p | $R^2$ | df |
| intercept | 4.0461 | 0.1902 | 21.27 | $<2x10^{-16}$ | 0.0216 | 1258 |
| age (at blood draw) | 0.0039 | 0.0024 | 1.608 | 0.1081 | | |
| bmi | 0.0170 | 0.0038 | 4.529 | $6.49x10^{-6}$ | | |
| sex | -0.0657 | 0.0381 | -1.723 | 0.0851 | | |
| **Model 2: linear regression between standardized residual and global ancestry** | | | | | | |
| Intercept | -0.0611 | 0.0810 | -0.754 | 0.451 | 0.0024 | 1258 |
| PNS | 0.0044 | 0.1436 | 0.031 | 0.976 | | |
| EAS | 0.1585 | 0.1125 | 1.409 | 0.159 | | |
| AFR | 1.2135 | 1.0259 | 1.183 | 0.237 | | |

Model 1 models the non-genetic covariates according to the heuristic described in the **Methods**. The residual from model 1 is then inverse normalized and tested in model 2.

**S4 Table: Details of the association statistics of the covariates and global ancestries of fasting insulin.**

| Model 1: linear regression between ln(insulin) and covariates | | | | | | |
|---|---|---|---|---|---|---|
| variables | estimate | std. error | T | p | $R^2$ | df |
| intercept | 0.4118 | 0.1721 | 2.393 | 0.0169 | 0.1585 | 1265 |
| age (at blood draw) | -0.0012 | 0.0022 | -0.565 | 0.5719 | | |
| bmi | 0.0523 | 0.0034 | 15.4 | $<2x10^{-16}$ | | |
| sex | 0.0751 | 0.0344 | 2.18 | 0.0294 | | |
| **Model 2: linear regression between standardized residual and global ancestry** | | | | | | |
| Intercept | -0.1160 | 0.0807 | -1.437 | 0.1508 | 0.0033 | 1265 |
| PNS | 0.2716 | 0.1433 | 1.896 | 0.0582 | | |
| EAS | 0.0138 | 0.1124 | 0.123 | 0.9021 | | |
| AFR | 0.7477 | 1.0244 | 0.73 | 0.4656 | | |

Model 1 models the non-genetic covariates according to the heuristic described in the **Methods**. The residual from model 1 is then inverse normalized and tested in model 2.

**S5 Table: Details of the association statistics of the covariates and global ancestries of HDL.**

| variables | | estimate | std. error | t | p | $R^2$ | df |
|---|---|---|---|---|---|---|---|
| Model 1: linear regression between HDL and covariates | | | | | | | |
| intercept | | 29.3271 | 2.9464 | 9.954 | $<2\times10^{-16}$ | | |
| age (at blood draw) | | 0.1310 | 0.0452 | 2.898 | 0.0038 | 0.0641 | 1700 |
| sex | | 7.6327 | 0.7329 | 10.415 | $<2x10^{-16}$ | | |
| Model 2: linear regression between standardized residual and global ancestry | | | | | | | |
| intercept | | 0.1489 | 0.0712 | 2.093 | 0.0365 | | |
| PNS | | -0.4715 | 0.1235 | -3.817 | $1.40x10^{-4}$ | 0.0169 | 1699 |
| EAS | | 0.1753 | 0.0967 | 1.813 | 0.0700 | | |
| AFR | | -1.4498 | 0.8777 | -1.652 | 0.0988 | | |
| Model 1A: linear mixed model between HDL and covariates, including nSES | | | | | | | |
| intercept | | 25.6 | 3.332 | 7.682 | $3.29\times10^{-14}$ | | |
| age (at blood draw) | | 0.1308 | 0.048 | 2.717 | 0.0067 | | |
| sex | | 7.665 | 0.770 | 9.96 | $<2\times10^{-16}$ | | |
| nSES | (Q2 vs. Q1) | 5.518 | 1.535 | 3.595 | $4.66\times10^{-4}$ | 0.0902 | 1555 |
| | (Q3 vs. Q1) | 2.89 | 1.474 | 1.96 | 0.0520 | | |
| | (Q4 vs. Q1) | 3.208 | 1.448 | 2.215 | 0.0287 | | |
| | (Q5 vs. Q1) | 5.922 | 1.394 | 4.248 | $4.37\times10^{-5}$ | | |
| Model 2A: linear regression between standardized residual, including nSES, and global ancestry | | | | | | | |
| intercept | | 0.0787 | 0.0749 | 1.051 | 0.2935 | | |
| PNS | | -0.3027 | 0.1301 | -2.327 | 0.0201 | 0.0091 | 1551 |
| EAS | | 0.1725 | 0.1014 | 1.701 | 0.0891 | | |
| AFR | | -0.9393 | 0.9099 | -1.032 | 0.3021 | | |

Model 1 models the non-genetic covariates according to the heuristic described in the **Methods**. The residual from model 1 is then inverse normalized and tested in model 2. Models 1A and 2A repeats the procedure but included quintiles of nSES levels in a mixed effect model (**Method**); in this case, the $R^2$ in Model 1A reported include both the fixed and the random effect.

**S6 Table: Details of the association statistics of the covariates and global ancestries of LDL.**

| variables | estimate | std. error | t | p | $R^2$ | df |
|---|---|---|---|---|---|---|
| Model 1: linear regression between LDL and covariates | | | | | | |
| intercept | 148.2936 | 7.1638 | 20.7 | $<2\times10^{-16}$ | 0.0045 | 1678 |
| age (at blood draw) | -0.0951 | 0.1098 | -0.866 | 0.3866 | | |
| sex | 4.6162 | 1.7807 | 2.592 | 0.0096 | | |
| Model 2: linear regression between rank-based inversed residual and global ancestry | | | | | | |
| intercept | -0.0434 | 0.0721 | -0.602 | 0.547 | $7.96\times10^{-4}$ | 1677 |
| PNS | 0.0736 | 0.1252 | 0.588 | 0.557 | | |
| EAS | 0.0720 | 0.0981 | 0.734 | 0.463 | | |
| AFR | -0.5192 | 0.8890 | -0.584 | 0.559 | | |

Model 1 models the non-genetic covariates according to the heuristic described in the **Methods**. The residual from model 1 is then inverse normalized and tested in model 2.

**S7 Table: Details of the association statistics of the covariates and global ancestries of TG.**

| Model 1: linear regression between ln(TG) and covariates | | | | | | |
|---|---|---|---|---|---|---|
| variables | estimate | std. error | t | p | $R^2$ | df |
| intercept | 4.7329 | 0.1044 | 45.32 | $<2\times10^{-16}$ | 0.0026 | 1691 |
| age (at blood draw) | 0.0002 | 0.0016 | 0.116 | 0.908 | | |
| sex | -0.0187 | 0.0256 | -0.732 | 0.4641 | | |
| edu4* | -0.0577 | 0.0293 | -1.969 | 0.0492 | | |
| Model 2: linear regression between rank-based inversed residual and global ancestry | | | | | | |
| intercept | -0.0972 | 0.0377 | -2.578 | 0.0100 | 0.0055 | 1691 |
| PNS | 0.1387 | 0.0654 | 2.119 | 0.0342 | | |
| EAS | 0.1426 | 0.0511 | 2.791 | 0.0053 | | |
| AFR | 0.1639 | 0.4635 | 0.354 | 0.7237 | | |

Model 1 models the non-genetic covariates according to the heuristic described in the **Methods**. The residual from model 1 is then inverse normalized and tested in model 2. * edu4 was a binary variable created from the original categorical variable of education status by grouping levels 1,2,3 and coded 0, while education status level 4 was coded as 1. This was done because there were no significant associations between education levels 1 through 3 and BMI.

**S8 Table: Details of the association statistics of the covariates and global ancestries of total cholesterol.**

| Model 1: linear regression between TC and covariates | | | | | | |
|---|---|---|---|---|---|---|
| variables | estimate | std. error | t | p | $R^2$ | df |
| intercept | 205.7 | 7.671 | 26.816 | $<2\times10^{-16}$ | 0.0190 | 1702 |
| age (at blood draw) | 0.0057 | 0.1176 | 0.048 | 0.961 | | |
| sex | 10.94 | 1.907 | 5.733 | $1.16\times10^{-8}$ | | |
| **Model 2: linear regression between rank-based inversed residual and global ancestry** | | | | | | |
| intercept | -0.0337 | 0.0716 | -0.471 | 0.6375 | 0.0046 | 1701 |
| PNS | -0.0441 | 0.1243 | -0.355 | 0.7228 | | |
| EAS | 0.2072 | 0.0972 | 2.132 | 0.0331 | | |
| AFR | -0.8084 | 0.8832 | -0.915 | 0.3602 | | |

Model 1 models the non-genetic covariates according to the heuristic described in the **Methods**. The residual from model 1 is then inverse normalized and tested in model 2.

**S9 Table: Details of the association statistics of the covariates and global ancestries of obesity.**

| Model 1: logistic regression based on covariates | | | | | | |
|---|---|---|---|---|---|---|
| variables | | estimate | std. error | z | p | df |
| intercept | | -0.9591 | 0.4966 | -1.932 | 0.0534 | 3083 |
| age (at baseline) | | -0.0139 | 0.0093 | -1.496 | 0.1346 | |
| t2d | | 4.1860 | 0.6733 | 6.217 | $5.06\times10^{-10}$ | |
| edu4* | | -0.2234 | 0.1079 | -2.07 | 0.0385 | |
| t2d:age | | -0.0526 | 0.0124 | -4.233 | $2.31\times10^{-5}$ | |
| Model 2: logistic regression between obesity and covariates | | | | | | |
| intercept | | -1.0136 | 0.5225 | -1.94 | 0.0524 | 3080 |
| PNS | | 1.2164 | 0.2349 | 5.178 | $2.24\times10^{-7}$ | |
| EAS | | -1.3596 | 0.2073 | -6.559 | $5.40\times10^{-11}$ | |
| AFR | | 1.3181 | 1.5591 | 0.845 | 0.3979 | |
| age (at baseline) | | -0.0170 | 0.0095 | -1.791 | 0.0734 | |
| t2d | | 4.4103 | 0.6901 | 6.391 | $1.65\times10^{-10}$ | |
| edu4* | | -0.0789 | 0.1113 | -0.709 | 0.4782 | |
| t2d:age | | -0.0562 | 0.0127 | -4.415 | $1.01\times10^{-5}$ | |
| Model 3: logistic mixed model including nSES | | | | | | |
| Intercept | | -1.6383 | 0.2000 | -8.193 | $2.55\times10^{-16}$ | 2827 |
| PNS | | 1.0774 | 0.2470 | 4.361 | $1.29\times10^{-5}$ | |
| EAS | | -1.3105 | 0.2146 | -6.105 | $1.03\times10^{-9}$ | |
| AFR | | 1.2973 | 1.5884 | 0.817 | 0.4141 | |
| age (baseline) | | -0.0196 | 0.0099 | -1.972 | 0.0487 | |
| t2d | | 1.2979 | 0.0978 | 13.275 | $<2\times10^{-16}$ | |
| edu4* | | -0.0317 | 0.1174 | -0.27 | 0.7872 | |
| nSES | (Q2 vs. Q1) | -0.1483 | 0.1664 | -0.891 | 0.3730 | |
| | (Q3 vs. Q1) | -0.3517 | 0.1648 | -2.134 | 0.0328 | |
| | (Q4 vs. Q1) | -0.1163 | 0.1602 | -0.726 | 0.4680 | |
| | (Q5 vs. Q1) | -0.4175 | 0.1598 | -2.613 | 0.0090 | |
| t2d:age | | -0.0523 | 0.0132 | -3.956 | $7.63\times10^{-5}$ | |

Model 1 models the non-genetic covariates according to the heuristic described in the **Methods**. Model 2 then includes global ancestries in addition to the significant covariates. * edu4 was a binary variable created from the original categorical variable of education status by grouping levels 1,2,3 and coded 0, while education status level 4 was coded as 1. This was done because there were no significant associations between education levels 1 through 3 and obesity. Model 3 included quintiles of nSES levels in a mixed effect model.

**S10 Table: Details of the association statistics of the covariates and global ancestries of Type-2 Diabetes.**

| | | estimate | std. error | z | p | Df |
|---|---|---|---|---|---|---|
| Model 1: logistic regression based on covariates | | | | | | |
| variables | | estimate | std. error | z | p | Df |
| intercept | | -7.4892 | 0.4224 | -17.729 | $<2\times10^{-16}$ | 3047 |
| age (at baseline) | | 0.0644 | 0.0054 | 11.814 | $<2\times10^{-16}$ | |
| bmi | | 0.1317 | 0.0079 | 16.754 | $<2\times10^{-16}$ | |
| edu3* | 3 vs (1 & 2) | -0.3022 | 0.0919 | -3.289 | 0.0010 | |
| | 4 vs (1 & 2) | -0.3631 | 0.1043 | -3.482 | $4.97\times10^{-4}$ | |
| Model 2: logistics regression between type 2 diabetes and covariates | | | | | | |
| intercept | | -8.4590 | 0.4556 | -18.566 | $<2\times10^{-16}$ | 3045 |
| PNS | | 0.8209 | 0.2179 | 3.767 | $1.65\times10^{-4}$ | |
| EAS | | 1.1765 | 0.1738 | 6.769 | $1.30\times10^{-11}$ | |
| AFR | | 0.3603 | 1.5188 | 0.237 | 0.8125 | |
| age (at baseline) | | 0.0660 | 0.0055 | 11.972 | $<2\times10^{-16}$ | |
| bmi | | 0.1384 | 0.0082 | 16.946 | $<2\times10^{-16}$ | |
| edu3* | 3 vs (1 & 2) | -0.3059 | 0.0930 | -3.289 | 0.0010 | |
| | 4 vs (1 & 2) | -0.3666 | 0.1066 | -3.44 | 0.0006 | |
| Model 3: logistic mixed model including nSES | | | | | | |
| Intercept | | -0.5990 | 0.1826 | -3.281 | 0.0010 | 2827 |
| PNS | | 0.7575 | 0.2271 | 3.336 | $8.51\times10^{-4}$ | |
| EAS | | 1.1569 | 0.1774 | 6.522 | $6.92\times10^{-11}$ | |
| AFR | | 0.2535 | 1.5109 | 0.168 | 0.8668 | |
| age (at baseline) | | 0.0625 | 0.0057 | 10.932 | $<2\times10^{-16}$ | |
| bmi | | 0.1326 | 0.0084 | 15.738 | $<2\times10^{-16}$ | |
| edu3* | 3 vs (1 & 2) | -0.2943 | 0.0958 | -3.073 | 0.0021 | |
| | 4 vs (1 & 2) | -0.3317 | 0.1110 | -2.987 | 0.0028 | |
| nSES | (Q2 vs. Q1) | -0.2271 | 0.1485 | -1.529 | 0.1262 | |
| | (Q3 vs. Q1) | -0.1418 | 0.1445 | -0.981 | 0.3265 | |
| | (Q4 vs. Q1) | -0.2876 | 0.1432 | -2.008 | 0.0447 | |
| | (Q5 vs. Q1) | -0.2808 | 0.1389 | -2.021 | 0.0433 | |

Model 1 models the non-genetic covariates according to the heuristic described in the **Methods**. Model 2 then includes global ancestries in addition to the significant covariates. Model 3 included quintiles of nSES levels in a mixed effect model. * edu3 was a ternary variable created from the original categorical variable of education status by grouping levels 1 and 2. This was done because there were no significant associations between education levels 1 and 2 with T2D.

**S11 Table: Details of the association statistics of the covariates and global ancestries of heart failure.**

| Model 1: logistics regression based on covariates | | | | | | |
|---|---|---|---|---|---|---|
| variables | | estimate | std. error | z | p | df |
| intercept | | -9.0826 | 0.5881 | -15.444 | $<2\times10^{-16}$ | 2218 |
| age (at baseline) | | 0.1012 | 0.0076 | 13.306 | $<2\times10^{-16}$ | |
| bmi | | 0.0928 | 0.0093 | 9.957 | $<2\times10^{-16}$ | |
| sex | | -0.4895 | 0.1063 | -4.607 | $4.09\times10^{-6}$ | |
| edu3* | 3 vs (1 & 2) | -0.2746 | 0.1210 | -2.269 | 0.0233 | |
| | 4 vs (1 & 2) | -0.3991 | 0.1426 | -2.798 | 0.0051 | |
| Model 2: logistics regression between heart failure and covariates | | | | | | |
| intercept | | -9.5113 | 0.6263 | -15.186 | $<2\times10^{-16}$ | 2215 |
| PNS | | 1.0465 | 0.2831 | 3.697 | $2.18\times10^{-4}$ | |
| EAS | | 0.2528 | 0.2338 | 1.081 | 0.2797 | |
| AFR | | 3.4653 | 1.8375 | 1.886 | 0.0593 | |
| age (at baseline) | | 0.1009 | 0.0077 | 13.195 | $<2\times10^{-16}$ | |
| bmi | | 0.0882 | 0.0096 | 9.189 | $<2\times10^{-16}$ | |
| sex | | -0.4909 | 0.1069 | -4.592 | $4.39\times10^{-6}$ | |
| edu3* | 3 vs (1 & 2) | -0.2432 | 0.1217 | -1.998 | 0.0457 | |
| | 4 vs (1 & 2) | -0.3227 | 0.1444 | -2.234 | 0.0255 | |
| Model 3: logistic mixed model including nSES | | | | | | |
| intercept | | -1.1414 | 0.2632 | -4.336 | $1.45\times10^{-5}$ | 2055 |
| PNS | | 1.0201 | 0.3024 | 3.374 | $7.41\times10^{-4}$ | |
| EAS | | 0.2775 | 0.2458 | 1.129 | 0.2588 | |
| AFR | | 3.4056 | 1.8889 | 1.803 | 0.0714 | |
| age (at baseline) | | 0.1018 | 0.0083 | 12.325 | $<2\times10^{-16}$ | |
| bmi | | 0.0899 | 0.0102 | 8.84 | $<2\times10^{-16}$ | |
| sex | | -0.5090 | 0.1120 | -4.546 | $5.46\times10^{-6}$ | |
| edu3* | 3 vs (1 & 2) | -0.2291 | 0.1282 | -1.788 | 0.0738 | |
| | 4 vs (1 & 2) | -0.3298 | 0.1532 | -2.153 | 0.0314 | |
| nSES | (Q2 vs. Q1) | -0.2241 | 0.2128 | -1.053 | 0.2923 | |
| | (Q3 vs. Q1) | -0.3938 | 0.2123 | -1.855 | 0.0636 | |
| | (Q4 vs. Q1) | -0.4033 | 0.2102 | -1.918 | 0.0551 | |
| | (Q5 vs. Q1) | -0.3256 | 0.2030 | -1.604 | 0.1087 | |

Model 1 models the non-genetic covariates according to the heuristic described in the **Methods**. Model 2 then includes global ancestries in addition to the significant covariates. Model 3 included quintiles of nSES levels in a mixed effect model. * edu3 was a ternary variable created from the original categorical variable of education status by grouping levels 1 and 2. This was done because there were no significant associations between education levels 1 and 2 with heart failure.

**S12 Table: Details of the association statistics of the covariates and global ancestries of hyperlipidemia.**

| variables | estimate | std. error | z | p | df |
|---|---|---|---|---|---|
| Model 1: logistics regression based on covariates | | | | | |
| intercept | -2.0533 | 0.3836 | -5.353 | $8.64\times10^{-8}$ | 2222 |
| age (at baseline) | 0.0528 | 0.0070 | 7.526 | $5.23\times10^{-14}$ | |
| edu4* | 0.2674 | 0.1144 | 2.338 | 0.0194 | |
| Model 2: logistics regression between hyperlipidemia and covariates | | | | | |
| intercept | -2.3231 | 0.4066 | -5.713 | $1.11\times10^{-8}$ | 2219 |
| PNS | 0.0652 | 0.2473 | 0.264 | 0.7920 | |
| EAS | 0.6973 | 0.2012 | 3.465 | $5.30\times10^{-4}$ | |
| AFR | 0.9841 | 1.7462 | 0.564 | 0.5731 | |
| age (at baseline) | 0.0535 | 0.0070 | 7.599 | $2.98\times10^{-14}$ | |
| edu4* | 0.2529 | 0.1158 | 2.184 | 0.0290 | |

Model 1 models the non-genetic covariates according to the heuristic described in the **Methods**. Model 2 then includes global ancestries in addition to the significant covariates. * edu4 was a binary variable created from the original categorical variable of education status by grouping levels 1,2,3 and coded 0, while education status level 4 was coded as 1. This was done because there were no significant associations between education levels 1 through 3 and hyperlipidemia.

**S13 Table: Details of the association statistics of the covariates and global ancestries of hypertension.**

| variables | estimate | std. error | z | p | df |
|---|---|---|---|---|---|
| Model 1: logistic regression based on covariates | | | | | |
| intercept | -5.2619 | 0.5108 | -10.302 | $<2\times10^{-16}$ | 2235 |
| age (at baseline) | 0.0859 | 0.0076 | 11.269 | $<2\times10^{-16}$ | |
| bmi | 0.0537 | 0.0092 | 5.812 | $6.17\times10^{-9}$ | |
| Model 2: logistics regression between hypertension and covariates | | | | | |
| intercept | -5.8251 | 0.5313 | -10.963 | $<2\times10^{-16}$ | 2232 |
| PNS | 0.3842 | 0.2570 | 1.495 | 0.135 | |
| EAS | 0.8783 | 0.2053 | 4.277 | $1.89\times10^{-5}$ | |
| AFR | -0.9459 | 1.7337 | -0.546 | 0.585 | |
| age (at baseline) | 0.0865 | 0.0077 | 11.296 | $<2\times10^{-16}$ | |
| bmi | 0.0584 | 0.0095 | 6.129 | $8.85\times10^{-10}$ | |

Model 1 models the non-genetic covariates according to the heuristic described in the **Methods**. Model 2 then includes global ancestries in addition to the significant covariates.

**S14 Table: Details of the association statistics of the covariates and global ancestries for ischemic heart disease.**

| Model 1: logistics regression based on covariates | | | | | | |
|---|---|---|---|---|---|---|
| variables | | estimate | std. error | z | p | df |
| intercept | | -6.5801 | 0.4983 | -13.204 | $<2\times10^{-16}$ | 2218 |
| age (at baseline) | | 0.0942 | 0.0068 | 13.862 | $<2\times10^{-16}$ | |
| bmi | | 0.0476 | 0.0083 | 5.764 | $8.22\times10^{-9}$ | |
| sex | | -0.4739 | 0.0932 | -5.087 | $3.63\times10^{-7}$ | |
| edu3* | 3 vs (1 & 2) | -0.2965 | 0.1066 | -2.782 | 0.0054 | |
| | 4 vs (1 & 2) | -0.3636 | 0.1221 | -2.979 | 0.0029 | |
| Model 2: logistics regression between ischemic heart diseases and covariates | | | | | | |
| intercept | | -6.7066 | 0.5205 | -12.886 | $<2\times10^{-16}$ | 2215 |
| PNS | | 0.2881 | 0.2481 | 1.161 | 0.2457 | |
| EAS | | 0.1445 | 0.1970 | 0.733 | 0.4633 | |
| AFR | | -0.7074 | 1.7560 | -0.403 | 0.6871 | |
| age (at baseline) | | 0.0939 | 0.0068 | 13.795 | $<2\times10^{-16}$ | |
| bmi | | 0.0471 | 0.0085 | 5.535 | $3.11\times10^{-8}$ | |
| sex | | -0.4685 | 0.0934 | -5.015 | $5.31\times10^{-7}$ | |
| edu3* | 3 vs (1 & 2) | -0.2894 | 0.1069 | -2.707 | 0.0068 | |
| | 4 vs (1 & 2) | -0.3473 | 0.1235 | -2.813 | 0.0049 | |

Model 1 models the non-genetic covariates according to the heuristic described in the **Methods**. Model 2 then includes global ancestries in addition to the significant covariates. * edu3 was a ternary variable created from the original categorical variable of education status by grouping levels 1 and 2. This was done because there were no significant associations between education levels 1 and 2 with ischemic heart disease.

**S15 Table: Details of the association statistics of the covariates and global ancestries for stroke and transient ischemic attacks.**

| Model 1: logistics regression based on covariates | | | | | |
|---|---|---|---|---|---|
| variables | estimate | std. error | z | p | df |
| intercept | -8.8502 | 0.6655 | -13.298 | $<2\times10^{-16}$ | 2235 |
| age (at baseline) | 0.1056 | 0.0087 | 12.084 | $<2\times10^{-16}$ | |
| bmi | 0.0344 | 0.0111 | 3.087 | 0.0020 | |
| Model 2: logistics regression between stroke and transient ischemic attacks and covariates | | | | | |
| intercept | -8.9589 | 0.7007 | -12.785 | $<2\times10^{-16}$ | 2232 |
| PNS | 0.3612 | 0.3370 | 1.072 | 0.2839 | |
| EAS | 0.0719 | 0.2738 | 0.263 | 0.7928 | |
| AFR | 2.1347 | 2.1583 | 0.989 | 0.3226 | |
| age (at baseline) | 0.1053 | 0.0088 | 11.991 | $<2\times10^{-16}$ | |
| bmi | 0.0320 | 0.0115 | 2.781 | 0.0054 | |

Model 1 models the non-genetic covariates according to the heuristic described in the **Methods**. Model 2 then includes global ancestries in addition to the significant covariates.

**S16 Table: Stratified analysis of association between global genetic ancestry and BMI among T2D cases and controls.**

Strata: Type 2 diabetes

Model 1: linear regression between BMI and covariates

| variables | estimate | std. error | t | p | $R^2$ | df |
|---|---|---|---|---|---|---|
| intercept | 44.3084 | 1.2509 | 35.421 | $<2\times10^{-16}$ | 0.0868 | 1290 |
| age (at baseline) | -0.2341 | 0.0216 | -10.859 | $<2\times10^{-16}$ | | |
| sex | 0.1986 | 0.3272 | 0.607 | 0.544 | | |
| edu4* | -1.0289 | 0.4105 | -2.506 | 0.0123 | | |

Model 2: linear regression between rank-based inversed residual and global ancestry

| variables | estimate | std. error | t | p | $R^2$ | df |
|---|---|---|---|---|---|---|
| Intercept | 0.1326 | 0.0946 | 1.402 | 0.161 | 0.0590 | 1290 |
| PNS | 0.2371 | 0.1492 | 1.588 | 0.112 | | |
| EAS | -0.8096 | 0.1196 | -6.769 | $1.96\times10^{-11}$ | | |
| AFR | 0.6460 | 1.0207 | 0.633 | 0.527 | | |

Strata: Non-type 2 diabetes

Model 1: linear regression between BMI and covariates

| variables | estimate | std. error | t | p | $R^2$ | df |
|---|---|---|---|---|---|---|
| intercept | 30.7277 | 0.9192 | 33.43 | $<2\times10^{-16}$ | 0.0251 | 1790 |
| age (at baseline) | -0.0420 | 0.0167 | -2.523 | 0.0117 | | |
| sex | -1.3266 | 0.2473 | -5.364 | $9.19\times10^{-8}$ | | |
| edu4* | -1.1036 | 0.2756 | -4.005 | $6.47\times10^{-5}$ | | |

Model 2: linear regression between rank-based inversed residual and global ancestry

| variables | estimate | std. error | t | p | $R^2$ | df |
|---|---|---|---|---|---|---|
| intercept | -0.1963 | 0.0664 | -2.959 | 0.0031 | 0.0730 | 1790 |
| PNS | 0.8745 | 0.1173 | 7.453 | $1.41\times10^{-13}$ | | |
| EAS | -0.5347 | 0.0917 | -5.832 | $6.48\times10^{-9}$ | | |
| AFR | 1.3586 | 0.8293 | 1.638 | 0.1016 | | |

Model testing was performed in the same manner as the global analysis with BMI (**S1 Table**), except for stratifying based on T2D disease status. Model column provided the final model with association coefficients. * edu4 was a binary variable created from the original categorical variable of education status by grouping levels 1,2,3 and coded 0, while education status level 4 was coded as 1. This was done because there were no significant associations between education levels 1 through 3 and BMI.

**S17 Table: Model of association between global ancestry and BMI, including interaction with type-2 diabetes.**

| Model 1: linear regression between BMI and covariates | | | | | | |
|---|---|---|---|---|---|---|
| variables | estimate | std. error | t | p | $R^2$ | df |
| intercept | 34.1237 | 0.7376 | 46.264 | $<2\times10^{-16}$ | 0.0212 | 3401 |
| age (at baseline) | -0.0832 | 0.0131 | -6.378 | $2.10\times10^{-10}$ | | |
| Sex | -0.7239 | 0.2006 | -3.608 | $3.13\times10^{-4}$ | | |
| edu4* | -1.2995 | 0.2347 | -5.538 | $3.29\times10^{-8}$ | | |
| Model 2: linear regression between rank-based inversed residual and global ancestry | | | | | | |
| intercept | -0.4586 | 0.0628 | -7.307 | $3.46\times10^{-13}$ | 0.1738 | 3080 |
| PNS | 0.8330 | 0.1110 | 7.506 | $7.95\times10^{-14}$ | | |
| EAS | -0.5164 | 0.0867 | -5.954 | $2.91\times10^{-9}$ | | |
| AFR | 1.1126 | 0.7844 | 1.418 | 0.1562 | | |
| t2d | 0.9937 | 0.1087 | 9.139 | $<2\times10^{-16}$ | | |
| PNS:t2d | -0.6338 | 0.1787 | -3.546 | 0.0004 | | |
| EAS:t2d | -0.1846 | 0.1419 | -1.301 | 0.1933 | | |
| AFR:t2d | -0.5000 | 1.2382 | -0.404 | 0.6864 | | |

Model 1 models the non-genetic covariates according to the heuristic described in the **Methods**, except for type-2 diabetes status. The residual from model 1 was then inverse normalized and tested in model 2, which includes global ancestries, type-2 diabetes status, and interactions between global ancestries and type-2 diabetes status. * edu4 was a binary variable created from the original categorical variable of education status by grouping levels 1,2,3 and coded 0, while education status level 4 was coded as 1. This was done because there were no significant associations between education levels 1 through 3 and BMI.

**S18 Table: Variants within the admixture signal region that were reported to be associated with the tested or related traits in GWAS catalog.**

| SNP ID | Chr | Pos (hg19) | Reported P-value | Associated Trait | Mapped Gene | Allele Frequencies | | | |
|---|---|---|---|---|---|---|---|---|---|
| | | | | | | MEC-NH | EUR | EAS | AFR |
| rs79976124 | 6 | 66618657 | $2x10^{-6}$ | type 2 diabetes | NUFIP1P, ADH5P4 | 0.198 (A) | 0.288 | 0.0784 | 0.0024 |
| rs10498828 | 6 | 65533066 | $9x10^{-6}$ | type 2 diabetes | EYS | 0.164 (T) | 0.218 | 0.109 | 0.128 |

Reported P-value, associated trait, and mapped genes were provided by the GWAS catalog [2]. Allele frequencies were either calculated from the imputed data of the 178 reference MEC Native Hawaiian individuals with estimated PNS ancestry > 90%, or obtained from 1000 Genomes Project. Frequencies were reported with respect to the minor allele in the Native Hawaiians, given in parenthesis next to the Native Hawaiian frequency estimates.

**S19 Table: Allele frequencies across populations for the most strongly associated variant in chr6 for T2D in single variant association test.**

| SNP ID | Chr | Pos (hg19) | Associated Trait | Derived Allele Frequencies | | | |
|---|---|---|---|---|---|---|---|
| | | | | MEC-NH | EUR | EAS | AFR |
| rs370140172 | 6 | 66205761 | T2D | 0.243 (C) | 0 | 0.008 | 0 |

Allele frequencies were either calculated from the imputed data of the 178 reference MEC Native Hawaiian individuals with estimated PNS ancestry > 90%, or obtained from 1000 Genomes Project (reported on dbSNP: https://www.ncbi.nlm.nih.gov/snp/). Frequencies were reported with respect to the derived allele, given in parenthesis next to the Native Hawaiian frequency estimates.

**S20 Table: Association results to T2D in 2,852 Samoan Replication Cohort.**

| rsID | Pos (hg19, chr6) | Effect Allele | Other Allele | EAF | BETA | SE | P-val (Samoa) | P-val (MEC-NH) |
|---|---|---|---|---|---|---|---|---|
| rs13213141 | 64295122 | A | G | 0.216 | -0.002 | 0.091 | 0.986 | $1.770 \times 10^{-4}$ |
| rs62415478 | 65472417 | C | G | 0.239 | 0.015 | 0.086 | 0.858 | $1.034 \times 10^{-4}$ |
| rs60268597 | 65473673 | A | G | 0.237 | 0.020 | 0.086 | 0.813 | $2.012 \times 10^{-4}$ |
| rs62415480 | 65475772 | T | C | 0.237 | 0.019 | 0.086 | 0.830 | $2.012 \times 10^{-4}$ |
| rs72648371 | 65507941 | T | C | 0.197 | 0.030 | 0.093 | 0.743 | $1.927 \times 10^{-4}$ |
| rs374288303 | 66156770 | C | T | 0.087 | 0.021 | 0.132 | 0.873 | $1.698 \times 10^{-5}$ |
| rs369186009 | 66197658 | C | G | NA | NA | NA | NA | $4.209 \times 10^{-5}$ |
| rs370140172 | 66205761 | C | T | 0.087 | 0.025 | 0.132 | 0.852 | $1.248 \times 10^{-5}$ |
| rs79261478 | 66348532 | T | C | 0.087 | 0.011 | 0.133 | 0.934 | $2.612 \times 10^{-5}$ |
| rs75566215 | 66358487 | C | T | 0.087 | 0.011 | 0.133 | 0.934 | $2.258 \times 10^{-5}$ |

We attempted to replicate the association of rs370140172 and nine other proxies showing the strongest single-variant associations with a cross-sectional population based study of Samoans recruited from Independent Samoa (**Methods**). EAF, effect allele frequency in Samoans. BETA and SE refers to the effect size and standard errors, respectively, from the logistic mixed model association tests in the Samoan cohort. P-val (Samoa) and P-val (MEC-NH) provide the p-value from the logistic mixed model association tests in the Samoan cohort and MEC Native Hawaiian cohort, respectively.

**S21 Table: Local ancestry inference using RFMix is robust to the choice of recombination map.**

| Genetic map 2 \ Genetic map 1 | | AA Map | | |
|---|---|---|---|---|
| | | CEU | NA | YRI |
| Default Map | CEU | 0.4629 | 0.0048 | 0.0013 |
| | NA | 0.0052 | 0.4803 | 0.0004 |
| | YRI | 0.0010 | 0.0005 | 0.0436 |

| Genetic map 2 \ Genetic map 1 | | Constant Map | | |
|---|---|---|---|---|
| | | CEU | NA | YRI |
| Default Map | CEU | 0.4603 | 0.0078 | 0.0009 |
| | NA | 0.0069 | 0.4787 | 0.0004 |
| | YRI | 0.0022 | 0.0008 | 0.0420 |

| Genetic map 2 \ Genetic map 1 | | AA Map | | |
|---|---|---|---|---|
| | | CEU | NA | YRI |
| Constant Map | CEU | 0.4605 | 0.0065 | 0.0023 |
| | NA | 0.0079 | 0.4785 | 0.0008 |
| | YRI | 0.0008 | 0.0005 | 0.0421 |

To evaluate the impact of recombination map on local ancestry inference, we used the 1000 Genomes AMR population. Following the same procedure used for Native Hawaiians, we identified through unsupervised ADMIXTURE analysis 49 Peruvian (PEL) and 3 Mexican (MEX) individuals from 1000 Genomes as having > 80% Native American ancestry. We then inferred local ancestry using RFMix in 71 HapMap3 MEX individuals using the constructed reference panel of 99 CEU, 108 YRI, and 52 NA individuals from 1000 Genomes. We used three recombination map in the local ancestry inference: a HapMap2 pooled recombination map, a mis-specified African-American map [3], and a constant map that assumes a constant rate of 1cM / Mb across the genome. We compared in pairwise fashion the concordance of inferred ancestry across common variants between runs, and calculated concordance rate as the sum of the diagonal of the contingency table. Across all comparisons, even when using a constant rate map, the concordance rate is extremely high (0.987, 0.981, and 0.981 for the comparisons of default vs. AA map, default to constant rate map, and constant rate to AA map, respectively), suggesting that the choice of recombination map does not strongly impact the local ancestry inference using RFMix.

**S22 Table: phenotype inclusion and transformation for metabolic and quantitative cardiovascular traits.**

| Trait | Inclusion criteria and transformation |
|---|---|
| Glucose | Remove individuals with non-fasting glucose, with measurement greater than 7 mmol/L, who are pregnant, or have type-2 diabetes. |
| Insulin | Remove individuals with non-fasting insulin, with measurement of glucose greater than 7 mmol/L, who are pregnant, or have type 2. Insulin is then log-transformed. |
| HDL | Medication adjustments were made per table below, based on the approach reported in PAGE consortium. If multiple medications were reported, only the correction factor with largest effect was applied. Exclude individuals who are pregnant or who had not fasted for at least 8 hrs before blood draw. |
| LDL | Medication adjustments were made per table below, based on the approach reported in PAGE consortium. If multiple medications were reported, only the correction factor with largest effect was applied. Exclude individuals who are pregnant or who had not fasted for at least 8 hrs before blood draw, or individuals with measured LDL greater than 400 mg/dL. |
| TG | Medication adjustments were made per table below, based on the approach reported in PAGE consortium. If multiple medications were reported, only the correction factor with largest effect was applied. Adjusted TG was then log-transformed. Exclude individuals who are pregnant or who had not fasted for at least 8 hrs before blood draw, or individuals with measured TG greater than 3000 mg/dL. |
| TC | Medication adjustments were made per table below, based on the approach reported in PAGE consortium. If multiple medications were reported, only the correction factor with largest effect was applied. Exclude individuals who are pregnant or who had not fasted for at least 8 hrs before blood draw. |
| Type-2 diabetes | Remove among diabetes cases those individuals who are pregnant or who have been diagnosed as type 1 diabetes. Remove cases with age under 20 and controls with glucose over 7 mmol/L. |

| Trait \ Medication | Fibrates | Statins | Bile acid sequestrants | Niacin | Cholesterol absorption inhibitors |
|---|---|---|---|---|---|
| HDL | -5.9 | -2.3 | -1.9 | -9.9 | 0 |
| LDL | +40.1 | +49.9 | +40.5 | +24.7 | +40.5 |
| TG | +57.1 | +18.4 | 0 | +89.4 | 0 |
| TC | +46.1 | +52.1 | 0 | +34.6 | +40.5 |

These traits were studied in PAGE consortium and we thus follow the same criteria and transformation.

**S23 Table: Descriptive summary statistics of the traits and covariates analyzed.**

Quantitative traits

| Traits | | Sample size | mean | s.d. |
|---|---|---|---|---|
| BMI/$m^2 \cdot kg^{-1}$ | | 3427 | 28.90 | 5.86 |
| Waist-to-hip ratio (WHR) | | 2699 | 0.91 | 0.0072 |
| Glucose/$mmol \cdot L^{-1}$ | | 1693 | 4.98 | 1.28 |
| Insulin/$pmol \cdot L^{-1}$ | | 1700 | 8.31 | 7.48 |
| HDL/$mg \cdot dL^{-1}$ | Before adjustment | 1715 | 42.35 | 15.55 |
| | After adjustment | 1711 | 41.61 | 15.60 |
| LDL/$mg \cdot dL^{-1}$ | Before adjustment | 1705 | 127.81 | 36.92 |
| | After adjustment | 1701 | 144.33 | 36.88 |
| TG/$mg \cdot dL^{-1}$ | Before adjustment | 1717 | 122.421 | 80.27 |
| | After adjustment | 1713 | 129.88 | 81.63 |
| TC/$mg \cdot dL^{-1}$ | Before adjustment | 1717 | 194.28 | 39.48 |
| | After adjustment | 1713 | 211.59 | 39.71 |

Categorical traits

| Traits | Total size | Level | |
|---|---|---|---|
| | | 0 = undiagnosed | 1 = diagnosed |
| T2D | 3109 | 1799 | 1310 |
| Obesity | 3427 | 2612 | 815 |
| HF | 2239 | 1683 | 556 |
| HYPERL | 2239 | 657 | 1582 |
| HYPERT | 2239 | 639 | 1600 |
| IHD | 2239 | 1351 | 888 |
| STROKE_TIA | 2239 | 1933 | 306 |

Quantitative covariates

| Covariate | Sample size | Mean | s.d. |
|---|---|---|---|
| Age at baseline | 3428 | 54.26 | 7.67 |
| Age at blood draw | 1717 | 64.08 | 8.09 |

Categorical covariates

| Covariate | Total size | Level | Number |
|---|---|---|---|
| Sex | 3428 | 1 = Male | 1512 |
| | | 2 = Female | 1916 |
| Education (edu) | 3406 | 1: ≤8th grade | 222 |
| | | 2: high school | 1190 |
| | | 3: some college or vocational school | 1177 |
| | | 4: college graduates | 817 |
| Cigs_per_day (cig) | 3381 | 1: 0 cigarettes smoked per day | 1483 |
| | | 2: ≤5 cigarettes smoked per day | 219 |
| | | 3: 6-10 cigarettes smoked per day | 429 |
| | | 4: 11-20 cigarettes smoked per day | 731 |
| | | 5: 21-30 cigarettes smoked per day | 353 |
| | | 6: > 31 cigarettes smoked per day | 166 |

Summary statistics reported after exclusion and transformation as described in S20 Table. For biomarkers (glucose, insulin, HDL, LDL, TG, and TC), a subset of participants were invited after cohort entry. Thus there is an age at baseline and an age at blood draw.
